# Single-cell epigenetic analysis reveals principles of chromatin states in H3.3-K27M gliomas

**DOI:** 10.1101/2021.11.02.466907

**Authors:** Nofar Harpaz, Tamir Mittelman, Olga Beresh, Ofir Griess, Noa Furth, Tomer-Meir Salame, Roni Oren, Liat Fellus-Alyagor, Alon Harmelin, Sanda Alexandrescu, Joana Graca Marques, Mariella G. Filbin, Guy Ron, Efrat Shema

**Author notes:** These authors contributed equally (Nofar Harpaz, Tamir Mittelman).

## Abstract

Cancer cells are highly heterogeneous at the transcriptional level and in their epigenetic state. Methods to study epigenetic heterogeneity are limited in throughput and information obtained per cell. Here, we adapted Cytometry by Time of Flight (CyTOF) to analyze a wide panel of histone modifications in primary tumor-derived lines of Diffused Intrinsic Pontine Glioma (DIPG). DIPG is a lethal glioma, driven by histone H3 lysine 27 mutation (H3-K27M). We identified two epigenetically distinct subpopulations in DIGP, reflecting inherent heterogeneity in expression of the mutant histone. These two subpopulations are robust across tumor lines derived from different patients and show differential proliferation capacity and expression of stem-cell and differentiation markers. Moreover, we demonstrate the use of this high-dimensional data to elucidate potential interactions between histone modifications and epigenetic alterations during the cell-cycle. Our work establishes new concepts for the analysis of epigenetic heterogeneity in cancer that could be applied to diverse biological systems.

## Introduction

Epigenetic regulation of genome function is fundamental for cellular differentiation, and its deregulation often promotes cancer (Flavahan et al., 2017; Valencia and Kadoch, 2019; Zhao and Shilatifard, 2019). Post-translational modifications (PTMs) of histones, which compose the nucleosome structure, regulate transcription by mediating interactions with the transcriptional machinery and by physically affecting DNA accessibility. In the last decade, single-cell RNA sequencing technologies have revolutionized our understanding of transcriptional heterogeneity within normal and malignant tissues (Baslan and Hicks, 2017; Suvà and Tirosh, 2019). Parallel techniques were developed to explore epigenetic cellular heterogeneity, mainly focusing on single-cell DNA methylation and chromatin accessibility (Cusanovich et al., 2015; Kelsey et al., 2017; Luo et al., 2018; Shema et al., 2019; Smallwood et al., 2014). Yet, histone PTMs are mainly analyzed by bulk methods such as Chromatin Immunoprecipitation and sequencing (ChIP-seq). Recently, single-cell ChIP-seq and Cut&Tag revealed the patterns of several histone PTMs in the mouse brain, during stem cell differentiation, and in breast cancer (Bartosovic et al., 2021; Grosselin et al., 2019; Rotem et al., 2015; Wu et al., 2021). Advances in optical microscopy and super-resolution imaging allowed single-cell spatial analysis of histone modifications (Boettiger et al., 2016; Woodworth et al., 2021; Xu et al., 2018). While these advances show great promise, they are limited in scale and throughput, allowing analysis of one or two modifications at a time per single cell. Recently, a single-cell approach based on CyTOF, was used to study the epigenome of blood immune cells, demonstrating epigenetic alterations occurring with age (Cheung et al., 2018), and following vaccination (Wimmers et al., 2021).

Mutations in epigenetic regulators and histone proteins are frequent in many types of cancer. One example is the lysine 27-to-methionine mutation of histone H3 (H3-K27M), which is the driving event for DIPG, a lethal pediatric cancer. More than 80% of DIPG cases show a H3-K27M mutation on one allele out of 32 that encode canonical histone H3.1 or the histone variant H3.3 (Filbin and Monje, 2019; Nacev et al., 2019; Phillips et al., 2020; Schwartzentruber et al., 2012; Wu et al., 2012). This gain-of-function mutation was shown to cause drastic alterations in the epigenome, in both *cis* (on the mutant histones themselves) and *trans* (epigenetic alterations to wild-type histones) (Furth et al., 2021; Zhang and Zhang, 2019). The most notable changes induced by H3-K27M are the global loss of the repressive modification tri-methylation of histone H3 on lysine 27 (H3K27me3), with concomitant gain in acetylation of that same residue (H3K27ac) (Bender et al., 2013; Brian et al., 2021; Chan et al., 2013; Lewis et al., 2013; Piunti et al., 2017).

Here, we propose a novel adaptation of CyTOF, with a unique custom-designed epigenetic panel, as a method to reveal epigenetic dynamics and heterogeneity in cancer. We demonstrate the power of this method in revealing: (1) Systematic effects of chromatin perturbations, such as the H3-K27M mutation in glioma; (2) Discovery of epigenetic heterogeneity within tumor lines, as exemplified by two distinct subpopulations in DIPG; (3) Identification of baseline correlations between chromatin modifications, suggesting mechanisms of co-regulation; and (4) Characterization of dynamic epigenetic processes in single cells. This high-dimensional single-cell data is instrumental in deducing principles of epigenetic heterogeneity and regulation in cancer.

## Results

### CyTOF Epi-Panel reveals global epigenetic alterations induced by the K27M-mutant histone

To explore CyTOF as a method to reveal systematic effects of chromatin perturbations in cancer, we examined glioma models expressing the mutant histone H3.3-K27M. We designed an epigenetics-oriented panel, containing antibodies targeting the H3-K27M histone mutant, 18 well-characterized histone modifications, 5 histones proteins (H1, H1.0, H3, H3.3 and H4) and 3 chromatin regulators. We extended this panel with oncogenic, cancer stem cell and glioma lineage markers, as well as cell-cycle specific indicators (Figure 1A). Each selected antibody was conjugated to a metal, taking into consideration several parameters in metal allocation to enable parallel reading of all the antibodies in the same cells and minimize technical noise (see Methods). Antibodies specificity was verified by introducing functional perturbations (Fig. S1A-D).

**Figure 1:**
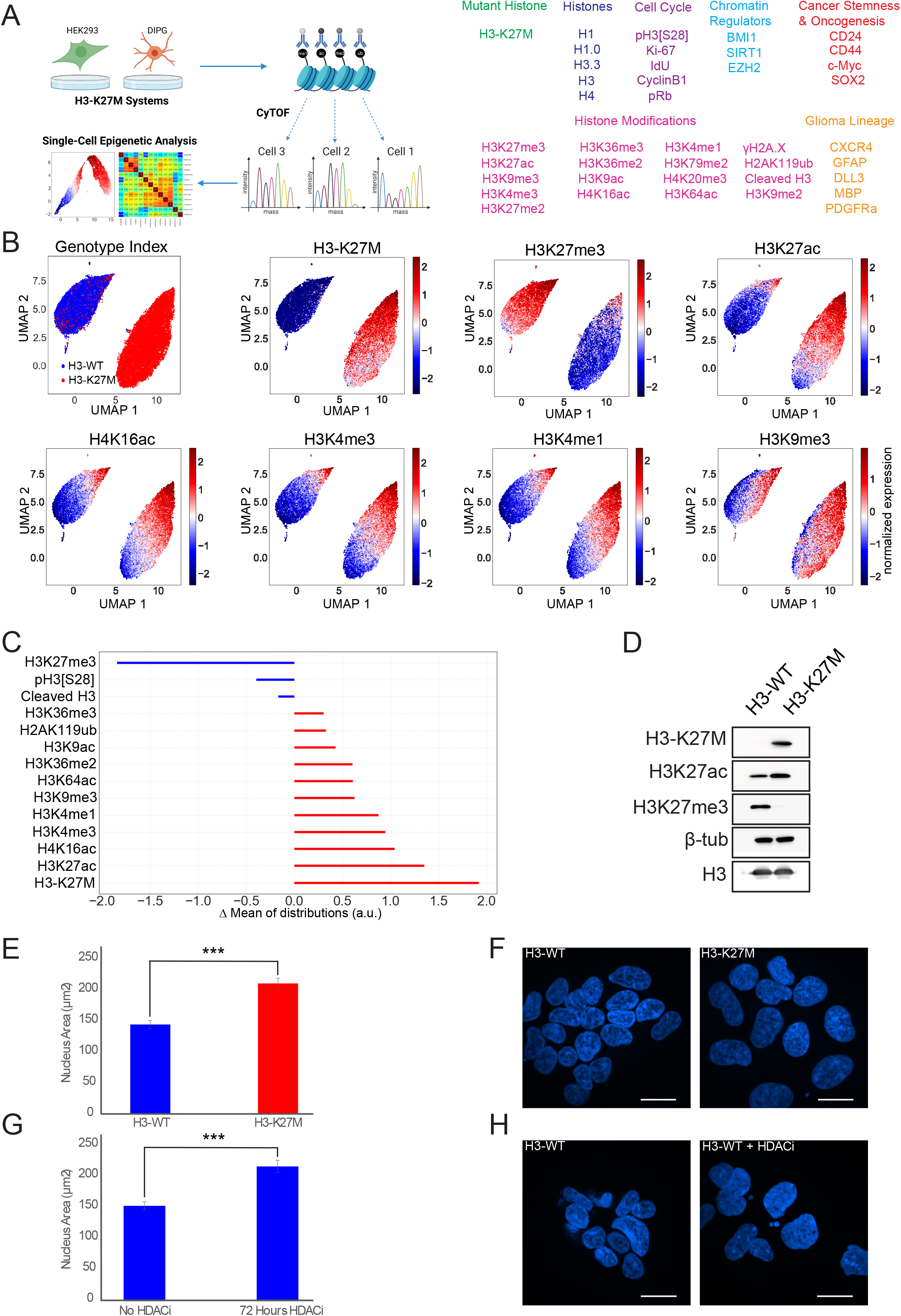
High-dimensional single-cell analysis of H3-K27M-induced epigenetic alterations. **(A)** Left: Scheme of CyTOF experimental setup and analysis for H3-K27M models. Right: Epigenetic-oriented antibody panel. The exact panel composition used for each experiment, and the number of cells analyzed, is described in Table S2 and Table S3, respectively. **(B-C)** HEK293 cells were induced for 10 days to express H3-WT or H3-K27M, and analyzed by CyTOF. Shown is one representative biological repeat out of three. **(B)** UMAP of CyTOF measurements after scaling and normalization. The H3-K27M signal was not included in the UMAP algorithm. Isogenic cells expressing H3-K27M are clustered separately from WT cells. **(C)** Fold change differences between the WT and H3-K27M expressing cells for the indicated epigenetic modifications. Mean values (after transformation, scaling and normalization) of the WT were subtracted from the mutant. (**D**) Western blot analysis of HEK293 cells induced as in (B-C) with the indicated antibodies. H3-K27M is robustly induced, leading to reduced levels of H3K27me3 concomitant with an increase in H3K27ac. Beta tubulin and H3 represent loading controls. (**E-F**) HEK293 cells were induced to express WT or H3-K27M for 72 hours, followed by DAPI staining to label nuclei. **(E)** Quantification of nuclei size (n=90, p-value = 3.2e-10), one-sided Student’s t-test was performed. Error bars represent SE. **(F)** Representative confocal images (scale = 20μm). (**G-H**) HEK293 cells were treated with the HDAC inhibitor (HDACi) Vorinostat for the indicated times, and stained with DAPI. **(G)** Nuclei size was calculated as in (E) (n=90, p-value = 1.3e-7). (**H**) Representative confocal images (scale = 20μm). Cells expressing H3-K27M or treated with HDACi have enlarged nuclei.

We first applied this epigenetic panel to an isogenic system of HEK293 cells with inducible expression of either wild type (WT) histone H3.3 or the H3.3-K27M-mutant. To minimize batch effect, samples were first barcoded so that subsequent processing and CyTOF runs could be done simultaneously. CyTOF experiments on three biological repeats showed very high reproducibility between experiments, validating the system (Figure 1B-C, S1E-F). We obtained single-cell measurements from 72,316 and 76,695 cells expressing the WT or K27M-mutant histone, respectively. Each measurement of histone modification was normalized to the level of core histones H3, H4, and H3.3 in that cell (Methods), followed by dimension reduction Uniform Manifold Approximation and Projection (UMAP) analysis (McInnes et al., 2018). Interestingly, HEK293 cells expressing the mutant H3.3 clustered separately from cells expressing the WT histone, using solely histone modifications for this algorithm and excluding the H3-K27M measurements (Figure 1B). This suggests that expression of H3-K27M results in global epigenetic alterations. Of note, we did detect a small fraction of the H3-K27M mutant cells that clustered with WT cells and indeed failed to express the H3-K27M construct, and were thus removed from further analysis (Figure S1G).

The most prominent epigenetic alterations induced by expression of H3-K27M were the global loss of H3K27me3 and gain in H3K27ac, as is well documented in the literature (Bender et al., 2013; Chan et al., 2013) (Figure 1B-C, S1F). In addition to these known effects, the extended panel revealed alterations in many other epigenetic modifications, associated with both active and repressed chromatin. Concomitant with the increase in H3K27ac, we observed a marked increase in lysine 16 acetylation on histone H4 (H4K16ac). The data also revealed a global increase in the activating marks mono-and tri-methylation of histone H3 on lysine 4 (H3K4me1, H3K4me3) as well as lysine 9 and 64 acetylations (H3K64ac, H3K9ac), albeit at a lower magnitude. These alterations suggest a more ‘open’ chromatin state induced by the H3-K27M mutation. However, there was also a parallel increase in the heterochromatin-associated mark tri-methylation of lysine 9 on histone H3 (H3K9me3), as was recently reported (Harutyunyan et al., 2020). In addition, we observed a reduction in the mitosis-associated phosphorylation of histone H3 (pH3S28), and a mild reduction in cleaved H3, a proteolytic form of histone H3 linked with mammalian differentiation (Duncan et al., 2008; Kim et al., 2016; Zhou et al., 2014) (Figure 1C). These results were highly reproducible across independent experiments with different H3-K27M induction times (4, 7 and 10 days) (Figure S1E-F), pointing to the robustness of the epigenetic alterations and the quantitative nature of CyTOF in revealing systematic effects mediated by this mutation.

The global increase in histone acetylations, suggesting a general opening of chromatin structure, motivated us to explore the shape of nuclei in cells expressing the mutant histone. Interestingly, we observed that HEK293 cells expressing H3-K27M had significantly larger nuclei than isogenic cells expressing WT H3.3 (Figure 1E-F). To examine whether the increase in nuclei size is indeed mediated by histone acetylations, we treated WT HEK293 cells with the histone deacetylase inhibitor Vorinostat (HDACi) for 72 hours. HDACi treatment led to an increase in nuclei size similar to that observed upon H3-K27M induction (Figure 1G-H). These results suggest that H3-K27M expression results in elevated histone acetylations, presumably leading to a more ‘open’ chromatin structure and enlarged nuclei.

### Single-cell profiling identifies epigenetic heterogeneity in H3-K27M tumor-derived primary cell lines

To examine H3-K27M-associated epigenetic alterations in a more biologically relevant system, we applied our CyTOF Epi-Panel to study two patient-derived DIPG cultures: SU-DIPG13, expressing the H3-K27M mutation endogenously, and SU-DIPG48, harboring WT H3. As expected, UMAP analysis of these lines, based solely on epigenetic marks, resulted in separate clustering of each patient (Figure 2A, S2A). H3-K27M expression was restricted to SU-DIPG13 cells, which also showed the expected low levels of H3K27me3 and high H3K27ac. Interestingly, while SU-DIPG48 cells clustered to a single center, SU-DIPG13 showed a clear division into two distinct subpopulations (Figure 2A-B, S2B). We repeated the UMAP analysis for SU-DIPG13 cells alone, and trained a gradient boosting algorithm (Chen and Guestrin, 2016) to determine the epigenetic features that affect assignment to the two clusters (Figure 2C, S2C-E). The results showed these two clusters mainly originated from higher expression of the H3-K27M mutant histone, H3K27ac and H3K4me1 pushing the cells towards one cluster, while increased levels of H3K9me3, cleaved H3 and pH3S28 pushed cells towards the second cluster. In fact, the algorithm allocated single cells into the correct cluster at 91% percent accuracy using reads from only four modifications: H3K27ac, cleaved H3, H3K9me3 and H3K4me1 (Figure S2E). Intriguingly, although DIPG cells all contain the H3-K27M mutation endogenously, the expression of the mutant gene varied in the population. This variability in H3-K27M expression likely induces epigenetic heterogeneity, generating the two distinct epigenetic subpopulations we termed H3-K27M-low and H3-K27M-high.

**Figure 2:**
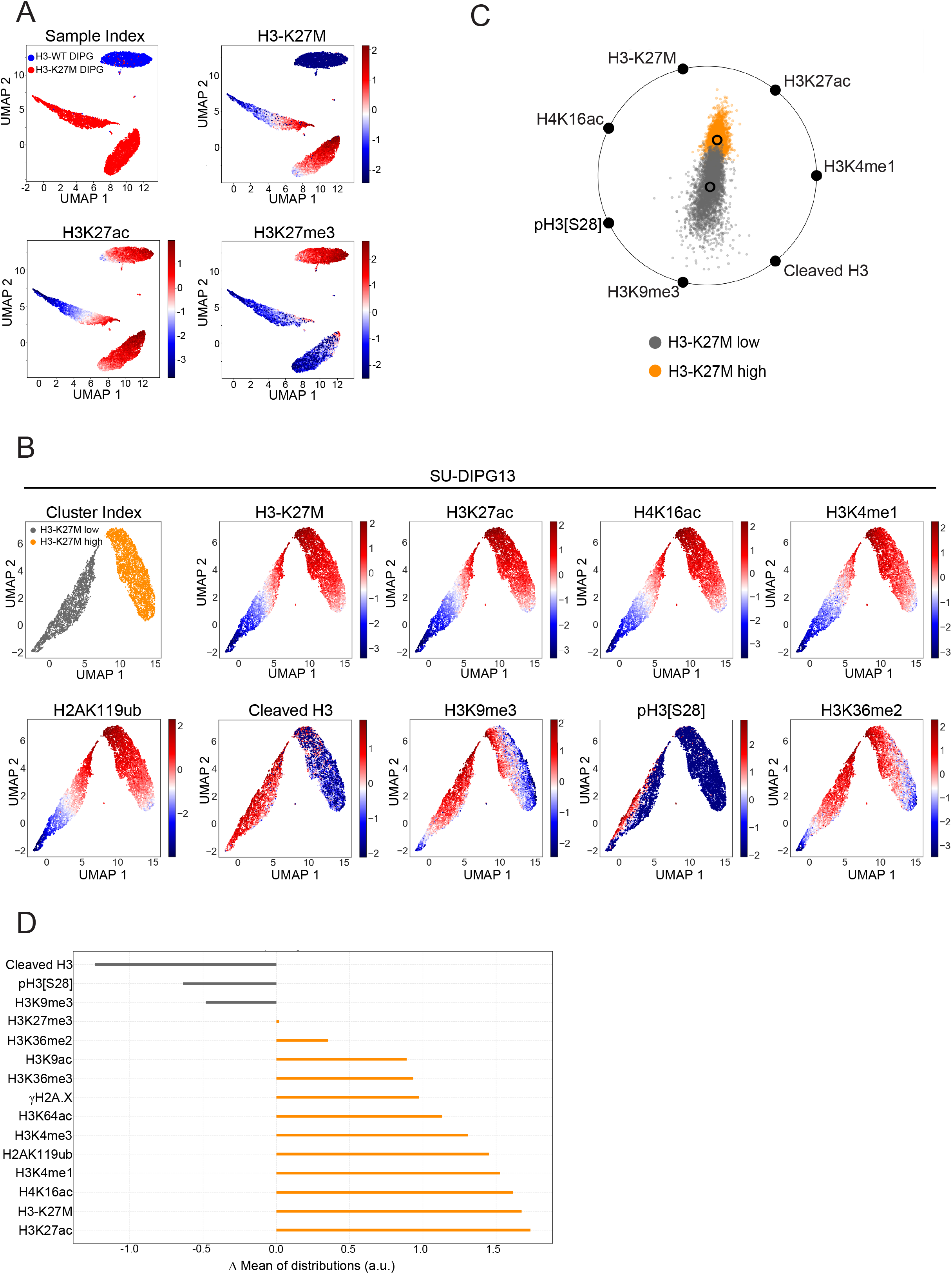
Two distinct epigenetic populations identified in H3-K27M tumor-derived cells. **(A)** SU-DIPG48 tumor-derived cells expressing WT H3 and SU-DIPG13 expressing H3-K27M were analyzed by CyTOF. UMAP based on all epigenetic marks measured shows separate clustering of the two tumor-derived lines (Top left: red/blue indicates the sample index). Also shown are the scaled, normalized levels of H3-K27M, H3K27ac and H3K27me3 in these cells. Shown is one representative biological repeat out of two. For panel composition and number of cells analyzed, see Tables S2-3. **(B)** UMAP of SU-DIPG13 cells, based on epigenetic marks as in (A), showing two distinct clusters. Levels of the indicated modifications in these cells are shown. (**C**) Spring plot (Sharko et al., 2008) representation of the gradient boosting algorithm, demonstrating the contribution of the top seven influencing epigenetic markers in the allocation of cells to clusters. (**D**) Fold change differences between the two clusters identified in SU-DIPG13 for the indicated epigenetic modifications. Mean values (after transformation, scaling and normalization) of the H3-K27M-low cluster were subtracted from the H3-K27M-high cluster.

To characterize this epigenetic heterogeneity, we examined fold change differences of all chromatin marks in these two subpopulations. Similar to the trends observed in the HEK293 system, the H3-K27M-high cluster showed elevation of all active modifications, mainly H3K27ac, H4K16ac and H3K4me1/3, and reduced levels of cleaved H3 and pH3S28 (Figure 2B-D, S2F). Interestingly, the H3-K27M-low cluster showed higher levels of the constitutive heterochromatin mark H3K9me3, which was upregulated by H3-K27M in the HEK293 cells (Figure 1C). These results support a complex regulation of this modification by H3-K27M: while the mutant histone induces an overall increase in this mark compared to WT H3.3 cells, it seems that lower expression of H3-K27M leads to higher H3K9me3 levels versus H3K9me3 levels in cells expressing high-H3-K27M. These results led us to explore the regulation of this modification by H3-K27M in a dynamic setup, as described below. Finally, we also applied our CyTOF panel to analyze two isogenic DIPG cell lines, where the H3.1 or H3.3-K27M alleles were knocked-out using CRISPR (Krug et al., 2019). Consistent with the results in the HEK293 isogenic system, loss of the mutant histone ‘rescued’ the most prominent epigenetic effects, reducing histone acetylations and increasing H3K27me3 and pH3S28 (Figure S2G-H).

### CyTOF reveals dynamic epigenetic alterations mediated by H3-K27M, suggesting distinct modes of regulation

To explore the dynamics of epigenetic alterations mediated by H3-K27M, we induced expression of the mutant histone in the HEK293 cells for 8, 16, 48 and 96 hours, followed by CyTOF with a focused panel of six epigenetic marks and H3-K27M. We observed a gradual increase in the levels of H3-K27M expression, reaching saturation at 48 hours (Figure 3A-C, S3A). A highly coordinated behavior was seen for H3K27ac, H4K16ac and H3K36me2. H3K27me3 levels, however, were not affected at the short induction, and started to decrease gradually only at later time points (Figure 3C). This is in line with the high stability of this modification, which is thought to be mainly ‘diluted’ during cell-cycle progression (Alabert et al., 2015; Rice and Allis, 2001). These results raise the intriguing possibility that the increase in histone acetylations occur prior to the loss of H3K27me3, and thus may be independent of H3K27me3 depletion. Alternatively, local loss of H3K27me3 might be sufficient to induce the acetylation phenotype, or alterations in other modifications not included in the panel.

**Figure 3:**
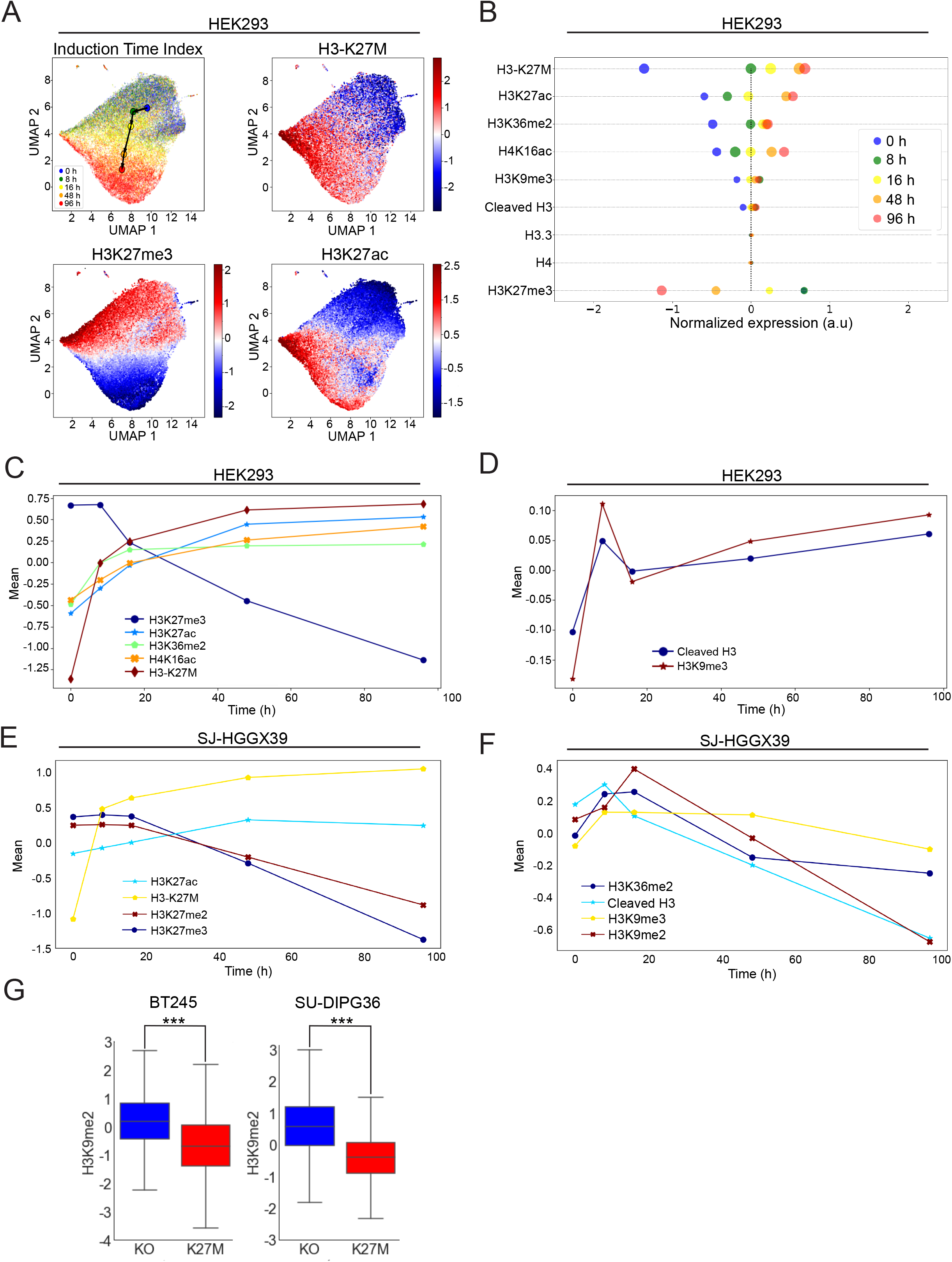
Dynamics of H3-K27M-induced epigenetic alterations. (**A-D**) HEK293 cells were induced to express H3-K27M for 8, 16, 48 and 96 hours, compared to cells expressing WT-H3 (marked as zero induction of H3-K27M), and analyzed by CyTOF with a panel of six epigenetic modifications, H3-K27M, and the core histones for normalization. (**A**) UMAP based on all epigenetic modifications measured shows spatial continuous separation of the cells by their time of induction. Also shown are the scaled, normalized levels of H3-K27M, H3K27ac and H3K27me3. (**B**) The mean of distribution of the indicated marks at the different time points. Dot size represents sample variance. (**C, D**) Mean values of the indicated modifications as a function of H3-K27M induction time. While the modifications in (C) show gradual increase/decrease with H3-K27M expression, cleaved H3 and H3K9me3 (D) are distinctly affected by differences in H3-K27M levels. To calculate the significance of changes over time, we performed Welch’s t-test for each modification at each of the time points, relative to the time point that preceded it. All changes are significant (p-value < 0.001) except for H3K27me3 levels at 0 to 8 hours, which remain the same. (**E, F**) SJ-HGGX39 cells were induced to express H3-K27M for 8, 16, 48 and 96 hours, compared to non-induced cells. CyTOF analysis was done as in (A), with the addition of H3K27me2 and H3K9me2 antibodies. Shown are the mean values of the indicated modifications as a function of H3-K27M induction time, as in (C, D). P-values were calculated as in (C, D): all changes are significant except for the change in H3K27me2 levels from 0 to 8, and 8 to 16 hours; the H3K36me2 change from 8 to 16 hours, and the H3K9me3 change from 8 to 16 hours, and 16 to 48 hours. The experiment was performed once with multiple time points. (**G**) H3K9me2 expression levels in SU-DIPG36 and BT245 cell lines, in the control (K27M) versus H3-K27M knock-out (KO) cells, as measured by CyTOF. P-values were calculated by Welch’s t-test. *** P-value < 0.001.

While most of the epigenetic marks showed a gradual unidirectional change (increase or decrease) over H3-K27M induction, for H3K9me3 and cleaved H3 we detected a complex non-monotonic behavior. At early time points, corresponding to low or moderate expression of H3-K27M, H3K9me3 and cleaved H3 expression spiked. At later time points, in which the expression of H3-K27M increased, H3K9me3 and cleaved H3 levels dropped, followed by a gradual, milder elevation (Figure 3D). At each of these points, H3K9me3 levels were higher than prior to induction, suggesting a general positive effect of H3-K27M on H3K9me3 levels. This result aligns with our data in HEK293 cells (Figure1C) and the SU-DIPG36 isogenic system (Figure S3B). Interestingly, the highest levels of H3K9me3 and cleaved H3 were detected in the presence of moderate, rather than high, expression of the mutant histone, in line with our observations that in SU-DIPG13, the subpopulation of H3-K27M-low cells contained higher levels of these marks compared to H3-K27M-high cells (Figure 2D). To examine this phenomenon in a more biologically relevant background, we established a similar system of inducible H3.3-K27M expression in the DIPG cells SJ-HGGX39, which contain WT H3.3. To expand our observations, we included antibodies targeting the di-methylations of H3 Lys9 and Lys27 (H3K9me2 and H3K27me2). Similar to the results in HEK293, expression of H3-K27M in these DIPG cells induced a gradual increase in H3K27ac, observed as early as eight hours following induction (Figure 3E, S3C-D). Of note, both the H3K27me2 and H3K27me3 modifications showed highly similar dynamics, decreasing only at later time points. Strikingly, the same non-monotonic behavior described above for H3K9me3 and cleaved H3 was seen also in these DIPG cells, with moderate rather than high H3-K27M expression associated with higher levels of these epigenetic marks (Figure 3F). Interestingly, H3K9me2 and H3K36me2 showed highly similar trends. While the di-and tri-methylation of histone H3 Lys9 shared a similar dynamic behavior, the overall levels of H3K9me2, as opposed to H3K9me3, decreased following H3-K27M expression. This result is in agreement with the DIPG isogenic systems, showing higher levels of H3K9me2 in cells knocked-out for the mutant histone (Figure 3G). Altogether, the data supports a complex relationship between H3-K27M expression and the histone modifications H3K9me2, H3K9me3, H3K36me2 and cleaved H3, which are highly dependent on the relative levels of H3-K27M expression. Moreover, these results provide further evidence for the importance of these epigenetic marks in dictating the two distinct epigenetic subpopulations in SU-DIPG13.

### The two epigenetic subpopulations are robust across different DIPG tumor-derived lines

Our findings of epigenetic heterogeneity in SU-DIPG13 prompted us to explore whether this phenomenon was unique to that cell line, or if it could be generalized to additional DIPG tumor-derived lines. To that end, we performed CyTOF in primary cell lines derived from three additional patients with H3-K27M mutation: SU-DIPG6, SU-DIPG25 and SU-DIPG38, as well as the WT H3 DIPG line discussed above, SJ-HGGX39. We also repeated the CyTOF for SU-DIPG13, aiming to examine more cells as well as expand our panel of antibodies (Table S2). UMAP analysis showed that all four H3-K27M DIPG cell lines presented two epigenetic clusters, although the proportion of cells in the H3-K27M-low cluster varied between cell lines and between experimental repeats of the same cell line (Figure 4A). Moreover, a shared UMAP of two mutant lines showed they clustered together, indicating a highly similar epigenetic behavior, unlike SU-DIPG48 and SJ-HGGX39 expressing WT H3, which clustered separately from the mutant lines (Figure 4B-C, 2A). Reproducing the previous CyTOF experiment on SU-DIPG13, in all mutant DIPG lines the H3-K27M-low cluster had higher mean levels of cleaved H3 and cells expressing pH3S28, as well as H3K9me2. The H3-K27M-high cluster was characterized by higher levels of H3K27ac, H4K16ac and H4K4me1/3 (Figure 4D-E, S4A-E). Overall, the epigenetic landscape of the two clusters was highly conserved and robust between the different cell lines and biological repeats (Figure 4F). The two epigenetic subpopulations were seen at all phases of the cell cycle, and thus do not reflect differences in cell-cycle stage (Figure S4F-I). Interestingly, we also observed an increase in EZH2, the catalytic unit of PRC2, in the H3-K27M-high cluster (Figure S4J). This result may reflect a feedback mechanism to the loss of H3K27me3, which needs to be further investigated.

**Figure 4:**
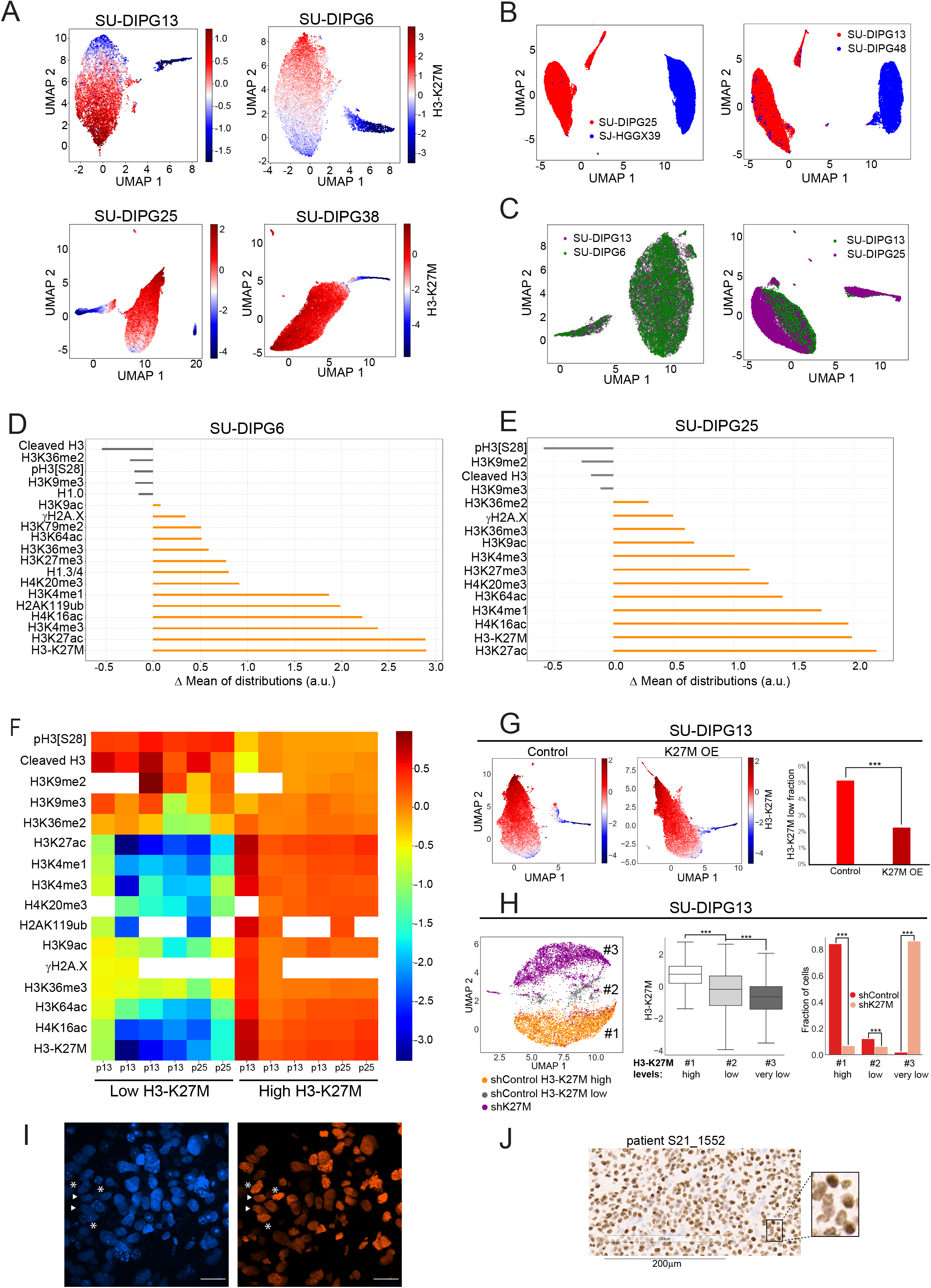
The two epigenetic states are robust across several patient-derived lines, with H3-K27M heterogeneity observed *in-vivo* in mouse model and in human tumors. (**A**) UMAP of four different patient-derived cell lines, based on all the epigenetic modifications measured, showing two distinct clusters. Levels of scaled, normalized H3-K27M are shown. Presented is one biological representative repeat out of four CyTOF experiments for SU-DIPG13, three for SU-DIPG25, one for SU-DIPG6 and one for SU-DIPG38. (**B**) UMAP of WT and H3-K27M patient-derived cell lines, based on all epigenetic modifications, analyzed together by CyTOF. Right: SU-DIPG13 (red) and SU-DIPG48 (blue). Left: SU-DIPG25 (red) and SJ-HGGX39 (blue). Colors represent samples’ barcodes, indicating that WT cells cluster separately from mutant cells. (**C**) UMAP of two H3-K27M DIPG cell lines analyzed together as in (B). Right: SU-DIPG13 (green) and SU-DIPG25 (purple). Left: SU-DIPG6 (green) and SU-DIPG13 (purple). Colors represent samples’ barcodes, indicating that each two lines, derived from different patients, are interchangeable, and that both show the two epigenetic clusters. (**D, E**) Representative fold change differences between the two clusters identified in SU-DIPG6 (**D**) and in SU-DIPG25 (**E**) for the indicated epigenetic modifications. Mean values (after transformation, scaling and normalization) of the H3-K27M-low cluster were subtracted from the H3-K27M-high cluster. (**F**) Heat map of the indicated modifications in H3-K27M-high and H3-K27M-low subpopulations, in four biological repeats of SU-DIPG13 and two of SU-DIPG25. (**G**) SU-DIPG13 cells were induced to express ectopic H3-K27M. Left: UMAP of non-induced cells (control) and cells following five days of induction (K27M OE-overexpression), based on epigenetic modifications as in (A). Shown are the levels of H3-K27M. Right: the percentage of cells allocated to the H3-K27M-low population in the control and following H3-K27M induction. P-values were calculated by Welch’s t-test. *** P-value < 0.001. (**H**) SU-DIPG13 cells were infected with shRNA targeting the H3F3A gene that carries the K27M mutation (‘shK27M’), or control shRNA targeting the H3F3B WT gene of H3.3 (‘shControl’), and analyzed by CyTOF. Left: joint UMAP analysis, based on all epigenetic modifications, of the shControl and shK27M cells. Colors represent samples’ barcodes; shControl cells were first barcoded as H3-K27M-high or H3-K27M low according to their allocation to these clusters in a UMAP performed on the control cells alone (shown in Figure S4K). Numbers (#1-3) represent areas with distinct H3-K27M expression levels, as presented in the middle panel. Right: The fraction of cells per sample allocated to areas #1-3. P-values were calculated as in (G). (**I**) SU-DIPG6 cells were injected to the pons of immunodeficient mice to form tumors. Representative confocal images (maximum projection) of DAPI (blue) and H3-K27M (red) staining. H3-K27M heterogeneity is observed: Asterisks mark cells expressing high levels of H3-K27M, and arrowheads mark cells expressing lower levels of the mutant histone, with similar DAPI signal. Scale bar = 20μm. (**J**) Human patient autopsy, stained for H3-K27M. Scale bar = 200μm.

To explore the causal effects of H3-K27M levels on the existence and proportion of cells in the two epigenetic subpopulations, we genetically manipulated SU-DIPG13 cells to express an inducible exogenous H3-K27M construct on top of its endogenous H3-K27M mutation. Following five days of H3-K27M induction, we detected a significant reduction in the fraction of cells consisting in the H3-K27M-low cluster (Figure 4G). Moreover, UMAP analysis revealed that the two epigenetic clusters were closer to each other in the H3-K27M-induced cells compared to the control SU-DIPG13, suggesting a higher degree of similarity in their epigenetic landscape. Next, we knocked down the endogenous mutant histone in SU-DIPG13 cells by infecting them with shRNA targeting the H3F3A gene that carries the K27M mutation (shK27M), or control shRNA targeting the H3F3B WT gene of H3.3 (shControl) (Silveira et al., 2019). UMAP analysis of shControl cells was done in order to define the H3-K27M-high and H3-K27M-low clusters (Figure S4K-L). Strikingly, a joint UMAP analysis of both the control and H3-K27M-depleted cells revealed three distinct areas that varied in their H3-K27M expression levels (Figure 4H). Cluster 1 corresponded to the H3-K27M-high subpopulation; while 84% of the shControl cells were allocated to this cluster, only 7% of the shK27M cells were associated with this area. Most of the shK27M cells (85.5%) showed strong depletion of the mutant histone and clustered separately from the H3-K27M-high and H3-K27M-low cells (Figure 4H, cluster 3). Interestingly, we identified 7.5% of the shK27M cells that expressed intermediate levels of H3-K27M, and clustered with the shControl H3-K27M-low cluster. This data indicates that in the shRNA system, H3-K27M-low cells expressed intermediate levels of the mutant histone, compared to the majority of shK27M cells that showed robust depletion of H3-K27M (thus showing greater similarity to H3-K27M knock-out cells). These intermediate H3-K27M levels generated a distinct epigenetic landscape, which was different from H3-K27M-high cells or cells completely depleted from the mutant histone, as evident from the UMAP analysis. Taken together, these results support a causal role for H3-K27M expression levels in dictating the two epigenetic subpopulations in DIPG patient derived cultures.

Finally, we aimed to examine whether this heterogeneity in H3-K27M expression could be detected in-vivo in mouse models and in patients diagnosed with H3-K27M gliomas. Thus, we established a DIPG orthotopic xenograft model by injections of SU-DIPG6 cells expressing GFP into the pons of immunodeficient mice (Grasso et al., 2015). H3-K27M immunofluorescence showed clear and robust heterogeneity in H3-K27M expression between tumor cells (Figure 4I). To better characterize this heterogeneity, we calculated the median intensity values of H3-K27M in 340-660 GFP-positive cells in the brains of each of three mice (Figure S4M-N). The histogram showed a broad distribution of intensity values, comparable to H3-K27M levels measured in the CyTOF experiments. Interestingly, while in the mouse brain the majority of cells seemed to have low H3-K27M expression, in the CyTOF experiment performed on cultured cells, we observed larger fraction of cells expressing high-H3-K27M (Figure S4M). These results may reflect differences in the microenvironment between cells grown in culture versus in-vivo growth in the brain. Importantly, immunohistochemical staining of H3-K27M in postmortem autopsy samples from three human patients also revealed intra-tumor H3-K27M heterogeneity in all specimens (Figure 4J, S4O). These results highlight the relevance of our findings to H3-K27M tumors in humans.

### The two epigenetic subpopulations show distinctive proliferation and differentiation features

To better characterize the two epigenetic subpopulations, we examined the expression patterns of stem cell, differentiation and cell cycle indicators included in our CyTOF panel. The histone variant H1.0 is known to be expressed heterogeneously within tumors, and H1.0 silencing by a fraction of tumor cells has been suggested to facilitate the transcription of oncogenes and stem-cell related genes (Torres et al., 2016). We found that H1.0 mean levels were reduced in the H3-K27M-high cluster, and a smaller fraction of cells expressed it in this cluster (Figure 5A-B, S5A). Moreover, H1.0 levels decreased in the HEK293 isogenic cells expressing H3-K27M compared to cells expressing WT H3.3, indicating the mutant histone affects H1.0 expression in diverse biological systems (Figure 5C). Interestingly, the cleaved form of H3, which is associated with cellular differentiation processes (Duncan et al., 2008; Kim et al., 2016; Zhou et al., 2014), also showed reduced levels in the H3-K27M-high cluster, suggesting a lower differentiation state of these cells (Figure 4D-E).

**Figure 5:**
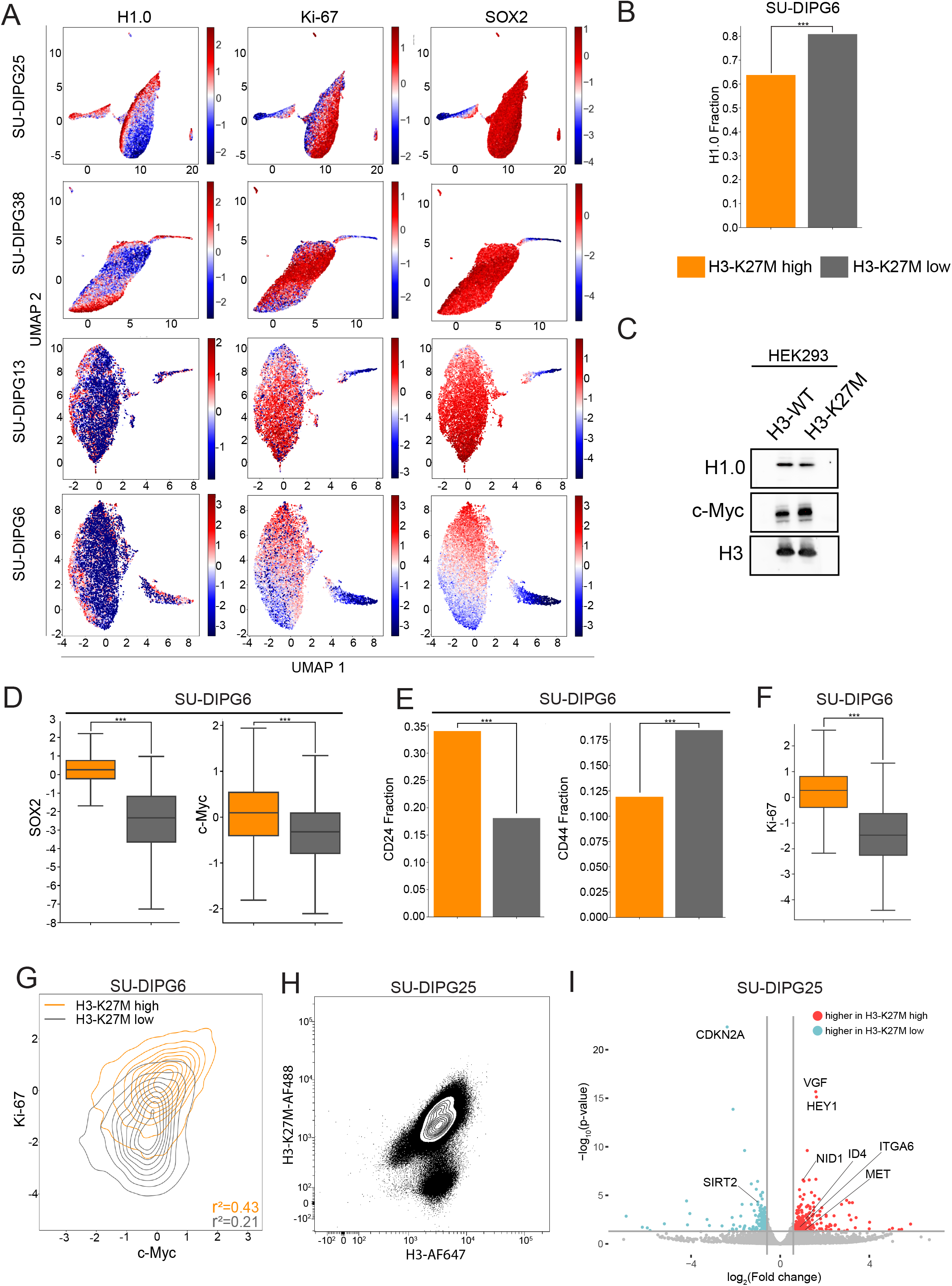
The two epigenetic subpopulations show distinct proliferation capacity and expression of oncogenic and cancer stem-cells markers. (**A**) UMAP analysis as in Figure 4A. Shown are the levels of H1.0, Ki-67 and SOX2. **(B)** Fraction of H1.0-positive cells in the two epigenetic subpopulations of SU-DIPG6. P-values were calculated by chi square test. *** P-value < 0.001. (**C**) Western blot analysis of HEK293 cells induced to express H3-WT or H3-K27M. H3-K27M expression leads to downregulation of H1.0 and upregulation of c-Myc. H3 represents loading control. (**D**) Box plots depicting CyTOF measurements of SOX2 and c-Myc (transformed and scaled) in SU-DIPG6 H3-K27M-high and H3-K27M-low clusters. P-values were calculated by Welch’s t-test. *** P-value < 0.001. **(E)** Fraction of cells positive for CD24 and CD44 in the two epigenetic subpopulations. P-values were calculated as in (B). (**F**) Box plots of Ki-67 expression levels in the two epigenetic subpopulations. P-values were calculated as in (D). (**G**) Contour plots of c-Myc versus Ki-67 in SU-DIPG6 H3-K27M-high (orange) and H3-K27M-low (gray) clusters. (**H**) Flow cytometry analysis of SU-DIPG25 cells, stained for H3-K27M-AF488 and H3-AF647. Cells were sorted based on H3-K27M levels for subsequent RNA-sequencing. Shown is one representative biological repeat out of three. (**I**) Volcano plot showing differentially expressed genes in the sorted SU-DIPG25 cells, expressing high versus low H3-K27M levels.

SOX2 and c-Myc are well-known drivers of the stem-cell state, as well as oncogenes in the context of gliomagenesis (Pajovic et al., 2020; Suvà et al., 2014). SOX2 was reported to be induced by the ectopic expression of H3-K27M in an oligodendrocyte progenitor cell line, and to play an essential role in inducing a stem-cell-like state in glioblastoma (Pajovic et al., 2020; Suvà et al., 2014). We observed elevated levels of both SOX2 and c-Myc in the H3-K27M-high cluster (Figure 5A,D, S5A). The c-Myc protein was also upregulated in the HEK293 cells expressing H3-K27M (Figure 5C, S5B). In addition, we examined markers for the neural progenitor cells (NPCs) and the mesenchymal-like cells (MES-like) CD24 and CD44, respectively (Neftel et al., 2019). Interestingly, we found enrichment of CD24+ cells in the H3-K27M-high cluster compared to the H3-K27M-low cluster, and an opposite trend was observed for CD44+ cells (Figure 5E, S5A). We also detected a mild increase in the levels of the oligodendrocytes progenitor cells (OPC) marker PDGFRa (Figure S5C). Other markers, including the glioblastoma lineage markers CXCR4, GFAP, DLL3 and MBP, and the chromatin regulators SIRT-1 and BMI1, did not behave differently between the clusters (Fig S4C-D). The elevated levels of SOX2, c-Myc, PDGFRa and CD24 in cells expressing high levels of H3-K27M, in addition to the reduced expression of H1.0 and cleaved H3, suggest that cells belonging to the H3-K27M-high cluster represent the fraction of tumor cells that had higher self-renewal, stem-cell-like properties.

To examine the proliferation capacity of the two subpopulations, we included in our CyTOF panel the proliferation marker Ki-67. In all four cell lines, the H3-K27M-high cluster expressed higher levels of Ki-67 when compared to the H3-K27M-low cluster (Figure 5A,F, S5A). Of note, despite reduced Ki-67 in the H3-K27M-low cells, a higher fraction of these cells were positive for the mitosis-associated pH3S28, perhaps indicating prolonged unproductive mitosis (pH3S28, Figure S4E). Interestingly, the single-cell analysis revealed H3-K27M-mediated dependency between Ki-67 and oncogenic c-Myc, as the correlation between these two markers increased significantly in the H3-K27M-high cluster (Figure 5G, S5D). These results support the notion that the less differentiated H3-K27M-high cells are more proliferative. Moreover, inclusion of cell-cycle indicators in the experiment revealed that while >98% of cells were actively cycling, the small fraction of cells that were found to be in G0 state (315 out of 18,977 cells in SU-DIPG25, and 167 out of 25,336 cells in SU-DIPG13) were exclusively allocated to the H3-K27M-low cluster. These results are in line with a recent single-cell RNA-seq study of DIPG tumors, showing that the majority of tumor cells are indeed less differentiated, and these are the cells with the higher proliferation capacity that presumably maintain the tumor’s aggressive behavior (Filbin et al., 2018).

To gain a better understanding of the differential transcription profiles of these two epigenetic subpopulations, we FACS-sorted SU-DIPG25 cells based on their expression levels of H3-K27M relative to H3, followed by RNA sequencing (Figure 5H-I). Differentially expressed genes in the H3-K27M-high and H3-K27M-low cells showed significant overlap with recently published gene signatures of DIPG xenograft cells expressing H3-K27M, versus cells depleted for the mutant histone by shRNA (Silveira et al., 2019). Among these genes that were both upregulated in H3-K27M-high cells and included in the published signature, were genes known to promote high proliferation and stemness, such as MET (Wallace et al., 2013), ID4 (Jeon et al., 2008), HEY-1 (Brun et al., 2018), VGF (Wang et al., 2018) and ITGA6 (Ying et al., 2014). The gene NID-1, whose expression levels correlated with shorter survival rates for patients (Zhang et al., 2021), was also elevated in H3-K27M-high cells. Finally, H3-K27M-low cells expressed higher levels of the tumor-suppressor gene CDKN2A (p16). Taken together, this analysis provides further support to the high proliferative state observed for H3-K27M-high cells. Interestingly, the histone deacetylase SIRT-2 was upregulated in the H3-K27M-low cells, perhaps contributing to the reduced levels of histone acetylations in this subpopulation.

### Single-cell CyTOF analysis uncovers potential co-regulation of histone modifications

We next aimed to explore whether the CyTOF single-cell multi-parametric data can be leveraged to study baseline correlations between histone modifications. Our underlying assumption is that histone modifications that are mechanistically linked (for example, deposited or removed by the same enzymes) should be highly correlated at the single-cell level. As we have measurements of multiple repressive and active modifications within the same cells, we might be able to deduce potential interactions between different epigenetic marks. We first calculated the pairwise correlations between all pairs of histone modifications associated with gene activation: H3K27ac, H3K9ac, H3K64ac, H4K16ac, H3K4me3 and H3K4me1. In both HEK293 cells and SU-DIPG48, expressing WT H3.3, H3 acetylations on lysine 9, 27 and 64 clustered with each other (Figure 6A-B and S6A-B). This result aligns with our hypothesis, as these modifications are known to be deposited by similar enzymatic complexes (EP300/CREBBP (Di Cerbo et al., 2014; Zhou et al., 2019)). Of note, H4K16ac, which is deposited by the distinct acetyltransferase Males absent On the First (MOF) (Smith et al., 2006), showed lower correlation and did not cluster with H3 acetylations, further supporting our hypothesis. Interestingly, we also observed a high correlation between H4K16ac and the H3K4me1 in both lines, and in HEK293 cells these modifications also correlated highly with H3K4me3 (Figure 6A-B, S6B). In HeLa cells, MOF was shown to form a stable complex with MLL1, and the purified complex had robust methyltransferase activity in depositing H3K4me1-3 (Dou et al., 2005). The high correlation between these modifications suggests this complex may be active in diverse cellular systems, generating synchronized deposition of these histone marks. In addition, recent work identified the PHD finger domain of MLL4, the major methyltransferase that deposits H3K4me1, as a specific reader of H4K16ac (Zhang et al., 2019).

**Figure 6:**
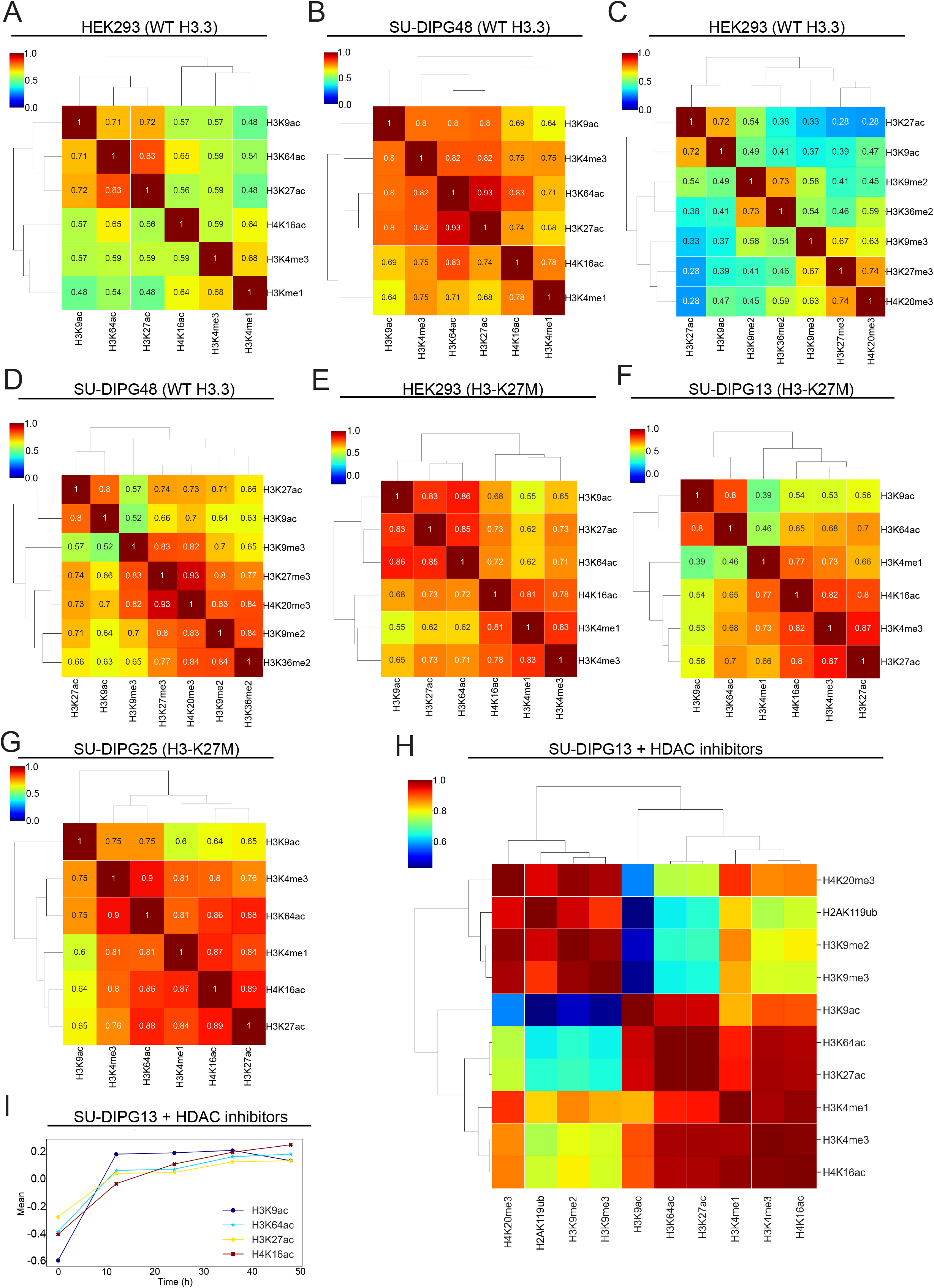
Single cell pairwise correlations suggest co-regulation between epigenetic marks. (**A-D**) Pearson pairwise correlation between histone modifications measured by CyTOF (after transformation, scaling and normalization) in WT HEK293 (A, C) and SU-DIPG48 (B, D) cells. **(A-B)** Coefficient of determination (R^2^) heat map matrix of active histone modifications. High correlations are observed between H3 acetylations on lysine residues 9, 27 and 64. H4K16ac is less correlated with H3 acetylations, and instead shows a high correlation with H3K4me1. (**C-D**) R^2^ heat map matrix of repressive histone modifications. H3K27me3 is highly correlated with H4K20me3. **(E)** R^2^ heat map matrix of active histone modifications measured in HEK293 cells expressing H3-K27M. (**F-G**) R^2^ heat map matrix of active histone modifications measured in (**F**) SU-DIPG13 and **(G)** SU-DIPG25 cells. (**H-I**) SU-DIPG13 cells were treated with the HDAC inhibitor Vorinostat for 0, 12, 24, 36 and 48 hours. Heat map represents the correlations between the means of distributions of the modifications at the different time points, indicating similarity in the trajectories of change of each modification in response to Vorinostat. High degree of correlation indicates modifications that varied together. (**I**) Mean values of the indicated histone acetylations in response to Vorinostat. To calculate the significance of changes over time, we performed Welch’s t-test for each modification at each of the time points, relative to the time point that preceded it. All changes are significant (p-value < 0.001) except for the following: the H3K64ac and H3K27ac changes from 12 to 24, and 36 to 48 hours, the H3K9ac change from 12 to 24, and 24 to 36 hours.

We next examined the correlations of repressive modifications with each other and with active chromatin marks (Figure 6C-D). The histone modifications H3K27me3 and H4K20me3, known to mark facultative heterochromatin, were highly correlated with each other, while H3K9me3, the hallmark of constitutive heterochromatin, showed significantly lower correlation with both of these marks. These observations are in line with the different genomic distributions of these modifications measured by ChIP-seq (Ernst et al., 2011) and immunofluorescence (Figure S6C). Interestingly, H3K9me2, but not H3K9me3, was highly correlated with the intergenic mark H3K36me2, perhaps reflecting their enrichment in similar genomic regions and the exclusion of H3K9me3 from H3K36me2 sites (Shirane et al., 2020; Weinberg et al., 2019). To examine whether the cell-cycle phase affects the correlations between the various histone marks, we repeated the correlation analysis for the sets of active and repressive modifications in cells found only at G1 or S phase (Figure S6D-G). While the repressive modifications showed similar clustering along the cell cycle, the active H4K16ac modification clustered with H3 acetylations during G1, yet in S phase it showed higher correlation with H3K4me3 and H3K4me1. This shift may imply a distinct regulation of H4K16ac during S phase. These results point to the potential of single-cell correlation analysis for uncovering co-regulation between epigenetic modifications.

To explore how epigenetic perturbations affect these patterns, we analyzed HEK293 cells expressing H3-K27M. We observed a general increase in all correlations between open chromatin marks in H3-K27M cells, in line with the global opening of the chromatin structure (Figure 6E). The most prominent increase in correlation was observed between H3K27ac and H4K16ac, the two modifications that were most upregulated by H3-K27M. To further generalize these observations, we explored pairwise correlations in SU-DIPG13, SU-DIPG25 and SU-DIPG6 tumor lines endogenously expressing H3-K27M (Figure 6F-G and 6SH). Similar to HEK293 cells expressing H3-K27M, in SU-DIPG13 and SU-DIPG25 cells H4K16ac clustered with H3K27ac, and not with H3K4me1 as in cells not expressing the mutant histone. This result reflects the epigenetic alterations caused by H3-K27M, coupling these two acetylations on histone H3 and H4.

To further explore the concept of histone PTMs crosstalk and provide additional validation to the correlations identified in single cells, we implemented a drug perturbation approach. We treated the SU-DIPG13 cells with the HDAC inhibitor Vorinostat, reported to have therapeutic potential for DIPG (Grasso et al., 2015), for several time points. This allowed us to calculate the trajectory of change for each modification, and examine the correlations between the trajectories of all modifications. Modifications that showed a similar trajectory of change in response to Vorinostat were clustered together (Figure 6H-I). Interestingly, pairs of modifications that were highly correlated at the single-cell level also showed a similar behavior upon Vorinostat treatment. These results provide additional support for a shared regulation of these modifications, and highlight the use of CyTOF data to deduce both baseline correlations between histone marks, as well as gain insights on how epigenetic perturbations affect various epigenetic pathways.

### Leveraging single-cell data to analyze dynamic chromatin alterations during the cell cycle

To further explore the utility of CyTOF in revealing dynamic processes, we aimed to leverage this data to more broadly identify global epigenetic alterations associated with different phases of the cell cycle. Thus, we combined our epigenetic CyTOF panel with the Maxpar cell cycle panel kit (Figure 1A), allowing us to define the cell cycle phase of each cell in the population. Analysis of the raw amounts of core histones revealed an increase in H3 and H4 levels when cells transitioned from G1 to S phase, in line with the expression of histone genes during DNA synthesis to enable formation of nucleosomes on the nascent DNA strand (Figure 7A, S7A). Accordingly, H3 and H4 levels further increased in G2 cells that contain double the amount of DNA, and dropped upon cell division (M-phase). To gain a robust view of the epigenetic changes associated with the cell cycle, we analyzed four different cell lines, including two lines that express H3-K27M. Across all these lines, we found that the chromatin in S-phase seems to be in a more ‘open’ state: it was enriched with active marks such as H3 acetylations, and showed reduced levels of the repressive H3K9me2/3 and H3K27me3 marks (Figure 7B-C, S7B-C). Interestingly, H4K16ac decreased during S phase, emphasizing the distinct regulation of H3 and H4 acetylations, as also indicated by their lower correlation during S-phase (Figure 7B-C, S7B-C, S6D-E). In contrast to the previously shown pattern of a global increase in histone methylations (Kheir and Lund, 2010), we observed a decrease in most methylations examined, including the transcription-associated mark H3K36me3. Unlike most methylations, H3K4me3 levels increased during S-phase, in agreement with the open chromatin state (Figure 7B-C, S7B-C). Of note, the elevation in histone acetylations during S-phase was milder in cells expressing H3-K27M, likely reflecting the high baseline levels of open chromatin marks in these cells. Finally, we also detected epigenetic alterations specific to mitosis, such as a significant elevation in γH2A.X (Figure 7D), a modification which is integral to the DNA damage response, and was previously shown to have an independent role during M phase (Ichijima et al., 2005)

**Figure 7:**
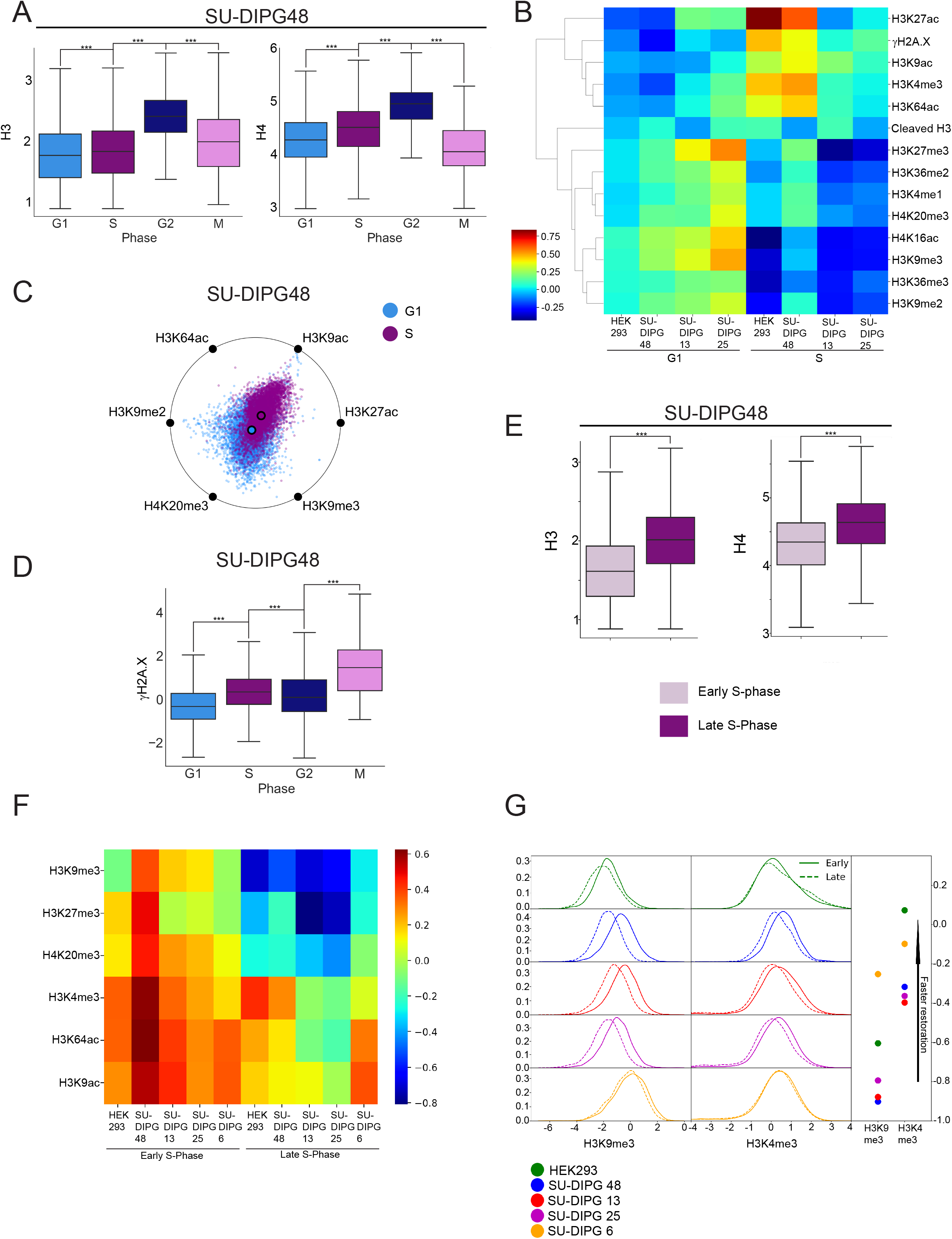
High-dimensional single-cell analysis of epigenetic alterations during cell cycle. (**A**) Box plots of the non-normalized scaled expression levels of H3 and H4 in different phases of the cell cycle, in SU-DIPG48 cells. P-values were calculated by Welch’s t-test. *** P-value < 0.001. (**B**) Heat map of the indicated histone modifications, scaled and normalized, in G1 and S phase cells, across four different cell lines: HEK293, SU-DIPG48, SU-DIPG13 and SU-DIPG25. A single repeat was done for each cell line. **(C)** Graphical representation of the epigenetic state of the G1 and S-phase populations of SU-DIPG48. (**D**) Box plots of scaled, normalized levels of γH2A.X in different phases of the cell cycle, in SU-DIPG48 cells. P-values were calculated as in (A). (**E**) Box plots of the non-normalized scaled expression levels of H3 and H4 during S-phase, indicating an increase in their expression levels. P-values were calculated as in (A). (**F**) Heat map of the indicated histone modifications, scaled and normalized, in early and late S phase, measured across five cell lines: HEK293, SU-DIPG48, SU-DIPG13, SU-DIPG25 and SU-DIPG6. A single repeat was done for each cell line. (**G**) Left: Histograms of H3K9me3 and H3K4me3 in early and late S phase in the five cell lines as in (F). Right: Dots represent the mean values of H3K9me3 and H3K4me3 modifications. P-values for the changes between early and late S phase, for each modification, were calculated by Welch’s t-test. All changes are significant with a p-value < 0.001, except for H3K4me3 in SU-DIPG6, for which the p-value = 0.0087.

Next, we leveraged our data to explore the epigenetic alterations that occur with time during S-phase. We used iridium labeling that marks total DNA content to divide the S-phase population accordingly: cells with high iridium are at a late stage of DNA synthesis, while cells with low iridium are in early S-phase. Similar to the increase observed in H3 and H4 levels between G1, S and G2 cells, we detected an increase in the levels of these core histones in late versus early DNA synthesis (Figure 7E). To explore the pattern of histone modifications along S-phase, we normalized their levels to the core histones H3 and H4, to account for the total increase in histones expression. As expected, almost all histone modifications showed decreased levels in late S-phase, compatible with the dilution of the parental modified histones and incorporation of new histones that are not yet modified (Figure 7F, S7D). Interestingly, the repressive histone modifications H3K27me3, H4K20me3 and H3K9me3 showed slower restoration in comparison to the active modifications H3K4me3, H3K64ac and H3K9ac. H3K9me3 levels, which were reported to be restored slowly and fully just by the next G1 cycle (Alabert et al., 2015), indeed showed a prominent decrease in late S-phase in all five cell lines examined (Figure 7F-G and S7D). The fold change of H3K4me3 in late versus early S-phase was lower than that of H3K9me3, indicating a faster restoration of this histone mark, in agreement with previous reports (Reverón-Gómez et al., 2018). Interestingly, among the modifications that showed very fast restoration across all cell lines were the acetylations of H3 on lysine 9 and 64. These results may reflect the more dynamic nature of histone acetylations versus methylations (Rice and Allis, 2001), and suggest faster kinetics of histone acetyltransferases versus methyltransferases. Moreover, studies of DNA replication timing showed that open, transcriptionally active chromatin is replicated early, while closed chromatin is replicated in late S-phase (Klein et al., 2021). This may account for the higher restoration of H3K9ac and H3K64ac seen in late S-phase, as well as H3K4me3, compared with close chromatin modifications such as H3K9me3.

## Discussion

In this study, we present the advantages of the CyTOF methodology to achieve a broad analysis of histone modification states in cancer cells at a single-cell resolution. The H3-K27M mutation in DIPG is known to drive drastic epigenetic alterations, however the extent of these alterations, the epigenetic marks that are influenced by it, and its heterogeneity within the population are unclear.

We identified downregulation of H3K9me2 and upregulation of H4K16ac as two of the most prominent effects of H3-K27M expression. Schachner et al. recently applied mass spectrometry to intact nucleosomes and showed high levels of H4K16ac on H3-K27M-mutant nucleosomes (Schachner et al., 2021), supporting a *cis* effect of the mutation on this specific H4 acetylation. The global increase in H4K16ac observed in our data strongly suggests a *trans* effect as well, similar to H3K27ac. Interestingly, our data suggests that these two acetylations, H3K27ac and H4K16ac, are likely to be distinctly regulated; first, they show a lower correlation between themselves compared to the correlations between H3 acetylations that are known to share enzymatic writers. Second, while H4K16ac is upregulated by H3-K27M, similarly to other acetylations, it shows a distinct behavior during S-phase, where its restoration to newly deposited histones is slow and does not follow H3 acetylations. Finally, studies report that H4K16ac is deposited by a specific writer, MOF, which does not target other lysine residues. Collectively, this data suggests unique functions for this acetylation that are not redundant with H3 acetylations.

The single-cell analysis revealed epigenetic heterogeneity within the DIPG tumor-derived cell lines, generating two distinct epigenetic subpopulations with unique characteristics. Interestingly, these epigenetic subpopulations are associated with inherent heterogeneity in H3-K27M expression, and were not detected in a DIPG line containing WT H3. While H3-K27M is expressed in a gradient across the population, the clear division into two clusters suggests a threshold effect for mediating various epigenetic alterations. In line with this notion is our finding that H3K9me2, H3K9me3, H3K36me2 and cleaved H3 levels are affected differently by the levels of H3-K27M in individual cells. Moreover, manipulating H3-K27M levels in DIPG cells by overexpression or depletion of the mutant histone induced a shift in the fraction of cells in these two subpopulations, indicating a causal role for H3K27M in generating this epigenetic heterogeneity. The heterogeneity in H3-K27M expression itself could also stem from epigenetic changes in its chromatin environment, generating a feedback mechanism that may further stabilize the two distinct states. Interestingly, while these two states are seen in four different patient-derived DIPG lines, and are reproducible across biological repeats, the fraction of cells in each cluster varies. This suggests that environmental conditions may affect the propensity of cells to be in each of the states. It is currently unclear how stable these subpopulations are, and whether cells can transition from one state to the other. Importantly, H3-K27M heterogeneity is also observed *in-vivo* in DIPG xenograft mouse model and in human tumors, supporting the relevance of these findings to the biology of this cancer.

Our work establishes new modes for the analysis of CyTOF data, focusing on epigenetic markers that are expressed in gradients in all cells rather than proteins with binary expression patterns that are often analyzed by CyTOF (i.e., cell surface markers indicative of cell identity, etc.). We developed a normalization strategy to account for technical variation that likely stems from differences in permeability between cells during the staining protocol, and is thus highly important when measuring nuclear proteins. We then leveraged the single-cell data to extrapolate on cross-talk between epigenetic marks. Interestingly, we identified a high correlation between H4K16ac and H3K4me3, which is supported by a reported interaction between the two writers that deposit these modifications: MOF and MLL1 (Dou et al., 2005). The high correlation suggests the deposition of these two marks may be coordinated in diverse biological systems. Finally, we show that the multiple measurements from individual cells can be leveraged to identify epigenetic changes associated with distinct cell states, for example phases of the cell cycle or early versus late DNA synthesis. Using the core histones for normalization allowed us to follow the restoration of histone modifications during S-phase, establishing the system as complementary to ChOR-seq (Reverón-Gómez et al., 2018). This approach can be applied to address diverse biological questions related to cancer and cellular differentiation.

## Limitations of the study

The CyTOF technology relies on the use of antibodies, and is thus susceptible to the general caveats associated with antibodies-based methodologies: non-specific binding, differences in avidity to nucleosomes that contain one or two copies of a modification, or differences in binding that stem from the presence or absence of other modifications on adjacent lysine residues. In the current study, we validated some of the antibodies by perturbation experiments (Figure S1A). Other antibodies were previously validated (Cheung et al., 2018; Fedyuk et al., 2021; Shema et al., 2016). Yet, we cannot rule out potential confounding effects of the use of antibodies on the data.

Another limitation of CyTOF is that it is restricted to the detection of global levels of modifications, and thus cannot determine the genomic localizations of the various marks. Moreover, minor changes in the global levels of some modifications might elicit significant downstream biological effects. Finally, changes in global levels measured by CyTOF might be influenced by the baseline abundance of the modification. For example, relatively small changes in highly abundant modifications may translate to large differences in terms of genomic occupancies. Nevertheless, our data indicates that the CyTOF technology is quantitative and highly sensitive even to minor changes, partially mitigating this limitation.

## Acknowledgments

We thank Y. Aylon and I. Tirosh for their important input. We are grateful to M. Monje for generously sharing with us the SU-DIPG48, SU-DIPG6, SU-DIPG6-GFP, SU-DIPG38, SU-DIPG25, and SU-DIPG13 cultures, and to N. Jabado for her kind gift of the isogenic BT245 and SU-DIPG36 cultures. We thank S. Baker, L. Kasper and the St. Jude Children’s Research Hospital Pediatric Brain Tumor Portal, https://pbtp.stjude.cloud, for the SJ-HGGX39 cell line and the sh^K27M^ constructs. We thank C. Raanan, M. Zerbib and I. Savchenko for their important contribution in establishing the DIPG mouse model. We thank G. Jona for his gracious assistance with the lentivirus system. The graphical abstract and Figure 1A were created using biorender.com. Funding: E.S. is an incumbent of the Lisa and Jeffrey Aronin Family Career Development chair. This research was supported by grants from the European Research Council (ERC801655), The Israeli Science Foundation (1881/19), The German-Israeli Foundation for Scientific Research and Development, the Israeli Council for Higher Education (CHE) via the Weizmann Data Science Research Center and Minerva.

## Author Contributions

N.H, T.M, G.R and E.S designed the study, contributed conceptually to the data analysis and wrote the manuscript. N.H, T.M and O.B established the antibodies panel and conducted the experiments. G.R and N.H performed CyTOF data analysis. T.S assisted in performing the CyTOF runs as well as designing the panel. N.F. and O.B. generated the HEK293 and SJ-HGGX39 inducible systems, and the SU-DIPG13 cell lines expressing H3-K27M or sh^K27M^. O.G assisted with RNA sequencing experiments. A.H, R.O and L.F. established the xenograft mice model and performed immunohistochemistry. S.A, J.M and M.F performed patients’ autopsy analysis.

## Declaration of interests

The authors declare no competing interests.

## STAR Methods

### Resource availability

#### Lead contact

Further information and requests for resources and reagents should be directed to and will be fulfilled by the lead contact, Efrat Shema.

#### Materials availability

Cell lines generated in this study are listed in the key resource table and are available upon request.

#### Data and code availability

Images of HEK293 nuclei, together with the tables detailing the analysis, have been deposited to Mendeley Data and are publicly available as of the date of publication. DOIs are listed in the key resources table. Images of mouse slides, together with the tables detailing the analysis, have been deposited to Mendeley Data and are publicly available as of the date of publication. DOIs are listed in the key resources table. CyTOF data has been deposited to Flow Repository and is publicly available as of the date of publication. DOIs are listed in the key resources table. MARS-Seq data has been deposited to GEO and is publicly available as of the date of publication. The accession number is listed in the key resources table.

All original code has been deposited at Zenodo and is publicly available as of the date of publication. DOIs are listed in the key resources table.

Any additional information required to reanalyze the data reported in this paper is available from the lead contact upon request.

### Experimental model and subject details

#### HEK293 and HEK293T cultures

HEK293 and HEK293T (female) cells were grown in complete DMEM containing 10% FBS (v/v) and 1% P/S (v/v) on 10cm dishes or 6-well plates and passaged by trypsination. Prior to doxycycline induction, G418 antibiotic was added to the medium at every passage at 0.5mg/ml to select against uninfected cells. Following induction, the medium was replaced daily with fresh medium containing 1μg/ml doxycycline. Prior to CyTOF experiments, HEK293 cultures were thoroughly triturated with a serological pipette after trypsination to ensure single-cell separation. For Immunofluorescence imaging, HEK293 cell were grown on glass coverslips (13mm, No.1) on 10cm dishes. Cells were grown in a standard humidified cell culture incubator at 37°C and 5% CO2. Cell cultures were regularly tested for mycoplasma contamination. HEK293 and HEK293T cells were not authenticated.

#### MCF7 cultures

MCF7 (female) cells for immunofluorescence imaging were grown in complete DMEM containing 10% FBS (v/v) and 1% P/S (v/v) on glass coverslips (13mm, No.1) on 10cm dishes and passaged by trypsination. Cells were grown in a standard humidified cell culture incubator at 37°C and 5% CO2. Cell cultures were regularly tested for mycoplasma contamination. MCF7 cells were not authenticated.

#### Glioma cultures

Glioma cells were grown according to a previously published protocol in T25 or T75 filter flasks in tumor stem medium (1X TSM) based on a 1:1 mixture of Neurobasal-A medium and DMEM-F12, and supplemented with: 10mM HEPES buffer, 1mM MEM sodium-pyruvate, 0.1mM MEM non-essential amino acids, 2mM L-alanyl-L-glutamine dipeptide, antibiotic antimycotic, 2% B-27 supplement w/o vitamin A (v/v), 20ng/ml H-EGF, 20ng/ml H-FGF-basic-154, 10ng/ml H-PDGF-AA, and 10ng/ml H-PDGF-BB and 2μg/ml heparin. When treated with Vorinostat, it was added to the growth medium to a final concentration of 1μM. Glioma cells were passaged as follows: The suspension containing the cells was collected into a tube and centrifuged at 300g for 5 minutes. In the meantime, 7ml of TrypLE Express added with DNase I (diluted 1:100 from a stock of 100mg/22.5ml) was added to the flask and incubate at 37°C to detach cells adhering to the bottom. After centrifugation, the supernatant conditioned medium was set aside in a new tube, and the trypsinated cells in the flask were added to the remaining pellet, resulting in 7ml of cells suspended in TrypLE Express. The flask was washed again with 5ml of TrypLE Express added with DNase 1 to collect all leftover cells and added to the trypsinated mixture, which was then triturated with a serological pipette and rotated for 10 minutes at 37°C to completely dissociate cells; this step was skipped for the adherent glioma lines: BT245, HGGX39 and SU-DIPG36. Following these 10 minutes, the cells were again triturated with a serological pipette, and the suspension then had 22ml of 1X HBSS added to it to dilute TrypLE express. The cells were then pelleted at 300g for 7 minutes and the supernatant was aspirated. Cells were resuspended in 1ml of HBSS and counted, and 1M were transferred to a T25 flask or 2M to a T75 flask. 2X TSM medium as described above was prepared in advance, and the cells were grown in a 1:1 mixture of 2X TSM and conditioned medium from the tube set aside. If glioma cells were harvested for CyTOF and not needed for maintenance, the conditioned medium was aspirated instead of kept, and cells were filtered through a 100μm mesh to remove any large aggregates after the addition of 22ml 1X HBSS. The sexes of the cells were as follows: SU-DIPG6 – female, SU-DIPG13 – female, SU-DIPG 25 – female, SU-DIPG – 36 – female, SU-DIPG 38 – female. Information about the sexes of the cell lines: SU-DIPG 48, BT245, and SJ-HGGX39 was not available. The SU-DIPG6, SU-DIPG13, SU-DIPG38, SU-DIP25, and SU-DIPG48 cells were a gift from M.Monje (Grasso et al., 2015). The SU-DIPG36 and BT245 KO cells and their parental K27M lines were a gift from N. Jabado (Krug et al., 2019). The SJ-HGGX39 cells were a gift from S. Baker. All cells were grown in a standard humidified cell culture incubator at 37°C and 5% CO2. Cell cultures were regularly tested for mycoplasma contamination. Glioma cells were not authenticated. Prior to doxycycline induction, G418 antibiotic was added to the medium at every passage at 0.25mg/ml to select against uninfected cells. Following induction, the medium was replaced daily with fresh medium containing 0.5μg/ml doxycycline. For shRNA experiments (detailed below), cells were grown with 0.5μg/ml puromycin.

#### NSG mice housing

Animal studies were approved by Institutional Animal Care and Use Committee (IACUC). NSG male mice 10-12 weeks old were housed and handled in a specific-pathogen-free, temperature-controlled (22°C ± 1°C) mouse facility on a 12/12 h light/dark cycle. Animals were fed a regular chow diet ad libitum.

#### Human subjects

S18-5613 is an 18 year-old female with a spinal tumor. S18-9426 is a 4 year-old male with a spinal nodule and leptomeningeal dissemination. S21-1552 is a 15 year-old male with a thalamic tumor. Since the IHC slides were acquired as part of clinical diagnosis, approval from an ethics committee was not required.

### Method details

#### Lentiviral transfection and selection

The pInducer20 H3.3 WT and H3.3-K27M plasmids were a gift from M. Suva. Lentiviral (3rd generation) packaging was performed by jetPEI-mediated transfection of HEK293T with appropriate plasmids (Tiscornia et al., 2006). Virus-containing supernatants were collected 48 hours following transfection, filtered, supplemented with 8µg/ml Polybrene, and added to target culture (HEK293, SJ-HGGX39 and SU-DIPG13). Infected HEK293 cells were selected with 0.5mg/ml G418, infected SJ-HGGX39 cells were selected with 0.25mg/ml G418 and infected SU-DIPG13 cells were selected with 0.2mg/ml G418. HEK293 cultures were induced with 1µg/ml doxycycline, while SJ-HGGX39 and SU-DIPG13 cultures were induced with 0.5µg/ml doxycycline. SU-DIPG13 cells used for shRNA experiments were selected with 0.5μg/ml puromycin. The shRNA plasmids targeting either H3F3A or H3F3B (pGIPZ-tRFP-miR-E-H3F3A or pGIPZ-tRFP-miR-E-H3F3B) were a gift from S. Baker. Lentiviral (2nd generation) production was performed by jetPEI-mediated transfection with Trans-Lentiviral Packaging plasmids (Wu et al., 2000). Virus-containing supernatants were collected 72h following transfection, filtered, supplemented with 8µg/ml polybrene, and added to SU-DIPG13 cells. Infected cells were selected with 0.5μg/ml puromycin for 12 days before CyTOF analysis.

#### Antibody metal conjugation

Antibodies were conjugated to metals using the Maxpar X8 Antibody Labeling Kit (Fluidigm) or the MIBItag Conjugation Kit (IONpath), according to the manufacturer’s protocols. Metals for conjugation were chosen in a way that would minimize noise and spillover between channels. This was done according to guidelines appearing in: (Han et al., 2018). Table S1 specifies the antibodies used and the metal allocated to each antibody.

#### CyTOF sample preparation

Sample preparation was similar to previously described protocols (Palii et al., 2019). Before harvesting, 5ml FACS tubes were coated with MaxPar Cell Staining Buffer (Fluidigm) for 30 minutes at RT or at 4°C overnight, to minimize loss of cells sticking to the sides of the tubes. Staining buffer was aspirated before starting the protocol. Up to 6M cells were harvested into 5ml FACS tubes, washed with in 4ml of MaxPar PBS (Fluidigm), then centrifuged at 400g for 10 minutes with slow break (used throughout the protocol). PBS supernatant was aspirated and the cells were labeled with 1.25μM Cell-ID Cisplatin (Fluidigm) in 1ml PBS for one minute for live/dead staining. Cisplatin was then quenched with 3ml DMEM+10% FBS pre-warmed to 37°C and the cells were centrifuged for 10 minutes at 400g. The supernatant was aspirated and cells were washed with 4ml of staining buffer, then centrifuged at 400g for 10 minutes. The supernatant was aspirated and the cells were then gently fixed in 1ml of nuclear antigen staining working solution (Maxpar Nuclear Antigen Staining Buffer Set, Fluidigm) for 30 minutes, then centrifuged at 700g for 10 minutes. The supernatant was aspirated and the cells were resuspended in 1ml nuclear antigen permeabilization buffer (Maxpar Nuclear Antigen Staining Buffer Set, Fluidigm), counted, and the supernatant was discarded as necessary so that no more than 3M cells would enter the next step. Then, the cells were centrifuged at 700g for 10 minutes, the supernatant was aspirated and the pellet was resuspended in 800μl nuclear antigen permeabilization buffer. Palladium-based barcodes (Cell-ID 20-Plex Pd Barcoding Kit, Fluidigm) were suspended in 100μl of nuclear antigen permeabilization buffer and immediately and completely mixed with the 800μl cells solution, then left on the bench top for 1 hour for complete barcoding. After 1 hour of barcoding, the cells were centrifuged at 700g for 10 minutes, then the supernatant was aspirated. The cells were then washed twice by resuspending them with 4ml nuclear antigen permeabilization buffer and centrifuging at 700g for 10 minutes. The cells were resuspended in 1ml nuclear antigen permeabilization buffer and counted. To ensure even and effective staining, a predetermined volume of antibody per 1M cells, and the total number of cells to be stained, had been titered, validated and prepared in advance, but within 4 hours of staining the cells. To minimize batch effect in sample preparation, the supernatant was discarded and samples were pooled into one tube so that all samples would contribute equally to staining and so that the total number of cells would not exceed the predetermined number, which was adjusted to be between 1M-2M cells per sample. The pooled sample was then centrifuged at 700g for 10 minutes. The total volume of antibodies to add was calculated, then the supernatant was aspirated and the cells were blocked in a volume of normal goat serum (NGS, Cell Signaling Technology) that would constitute 10% of the final solution containing the antibodies. After 10 minutes of blocking, antibodies were added to the solution and the solution was left on the bench top for 30 minutes for staining. Cells were then washed twice by suspending them in 4ml of staining buffer and centrifuging at 700g for 10 minutes. The supernatant was aspirated and around 50μl of supernatant residual volume was left and in which the cells were resuspended. The cells were fixed in 1-2ml 10% formalin by slowly adding it to the suspension while intermittently tapping the tube. The cells undergoing fixation were kept at 4°C overnight with gentle rocking. The next day, the formalin solution was supplemented with Cell-ID Intercalator-Iridium (Fluidigm) to achieve a final concentration of 125nM to label the DNA of the cells. The cells were then washed twice by suspending them in 3-4ml staining buffer and centrifuging at 700g for 10 minutes. The supernatant was aspirated and the cells were washed twice by resuspending them in Maxpar Cell Acquisition Solution (Fluidigm) and centrifuging at 700g for 10 minutes. Cells were resuspended in 1ml of cell acquisition solution containing 1:10 dilution of EQ Four Element Calibration Beads (Fluidigm), counted, and adjusted to attain a concentration of about 3M cells/ml. Cells were filtered through a 35μm mesh cell strainer (Falcon) before acquiring data on a Fluidigm Helios CyTOF system.

#### CyTOF data analysis

##### Data processing

CyTOF data underwent the following pre-processing prior to analyses: First, the CyTOF software by Fluidigm was used for normalization and concatenation of the acquired data. Then, several gates were applied using the Cytobank platform (Beckman Coulter): First, the normalization beads were gated out using the 140Ce channel. Then, live single cells were gated using the cisplatin 195Pt, iridium DNA label in 193Ir, event length, and the Gaussian parameters of width, center, offset and residual channels. CyTOF software was then used for samples de-barcoding.

##### Data manipulation and scaling

Before beginning the analysis procedure, the cells were gated using the core histones (H3, H3.3, and H4). The gate allowed only cells with a minimum raw value of 5 for all core histones into the next steps of the analysis. The gating on the core histones was followed by a gate on a subset of the epigenetic markers (H3K36me3, H3K4me3, H3K36me2, H4K16ac, H2Aub, H3K4me1, H3K64ac, H3K27ac, H3K9ac, H3K27me3, and H3K9me3), gating for a minimum raw value of 2 for said modifications. To deal with the different sensitivity of the CyTOF apparatus to the various markers a hyperbolic arcsine transform (with a scale factor of 5) was first applied to the data, followed by a Z-transform:

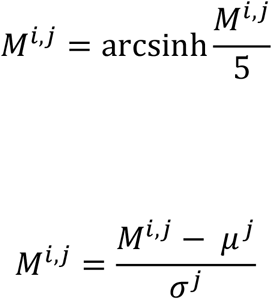

Where i denotes the observation, j denotes the column (modification), *µ*^*j*^ is the mean of column j, and *σ*^*j*^ is its standard deviation. Z-transform scaling was done at the same time on samples that were acquired together, such as the HEK293 WT and H3-K27M cell lines.

##### Epigenetic markers normalization

Based on the assumption that the core histones H3, H3.3, and H4 should be expressed at a similar level in all cells, systematic effects were removed from the measurement by subtracting a linear combination of the three core histones from the epigenetic values of all Observations 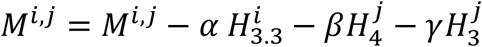,

Where the coefficients *α, β*, and *γ* are chosen by a least squares minimization procedure to minimize the sum of the variances of the core histones across all observations: *σ*(*H*_3.3_) +*σ*(*H*_4_) + *σ*(*H*_3_). A reduction of a few dozen percent in the sum of variances was observed after the subtraction, indicating the removal of systematic effects.

#### FACS staining and sorting

Before harvesting, 5ml FACS tubes were coated with wash buffer (2mM EDTA, 0.5% BSA (w/v) in PBS) for 30 minutes at RT or at 4°C overnight, to minimize loss of cells sticking to the sides of the tubes. For all buffers, RNasin Plus Ribonuclease Inhibitor was added to a concentration of 0.4U/μl, and the entire protocol was carried out on ice and at 4°C. Up to 10M cells were harvested and washed in 1ml of PBS and centrifuged at 400g for 5 minutes. The supernatant was aspirated and the cells were suspended in a 100μl PBS solution containing 1:400 Zombie Violet Fixable Viability Dye for 30 minutes for live/dead staining. The dye was quenched by adding 900μl of blocking buffer (2mM EDTA, 10% BSA (w/v) in PBS) and the cells were centrifuged at 400g for 5 minutes. The supernatant was aspirated and the cells were washed in 1ml PBS and centrifuged at 400g for 5 minutes. The cells were suspended in 100μl PBS, then had methanol pre-chilled to -20°C *slowly* added to them while tapping the tube, then kept on ice for 10 minutes for fixation and permeabilization. The cells were centrifuged at 900g for 5 minutes and the supernatant was aspirated. The cells were washed twice by suspending them in 1ml wash buffer and centrifuging at 900g for 5 minutes. The supernatant was aspirated and the cells were suspended in 100μl saturated ammonium sulfate solution containing 0.8U/μl RNasin Plus Ribonuclease Inhibitor and kept on ice for 10 minutes, and were then washed twice with 1ml was buffer and centrifuged at 900g for 5 minutes. The supernatant was aspirated and the cells were suspended in 100μl blocking buffer and kept on ice for 10 minutes, then primary antibodies (α-H3-K27M-AF488 and α-H3-AF647) were added to a final dilution of 1:500 for 30 minutes of staining. The cells were washed with 1ml wash buffer, then centrifuged at 900g for five minutes. The supernatant was aspirated and the cells were suspended in 1ml FACS buffer (0.1mM EDTA, 0.5% BSA (w/v) in PBS) and strained through a 40μm mesh. Flow cytometry analysis and sorting were performed on a FACSAria III instrument (BD Biosciences) equipped with a 407, 488, 561 and 633 nm lasers, using a 100 um nozzle, controlled by the BD FACSDiva software v8.0.1 (BD Biosciences).

#### RNA extraction and MARS-Seq

RNA was extracted from samples using a Qiagen RNeasy Micro Kit (Qiagen #74004) following the manufacturer protocol and stored at -80°C until needed. Library preparation for MARS-Seq from extracted RNA was done according to a published protocol using a NextSeq 500/550 High Output Kit v2.5 and sequencing was done on a NextSeq500 machine (Jaitin et al., 2014).

#### Western blot analysis

For western blot analysis, cell pellets were resuspended in Laemmli sample buffer containing 50mM DTT and vortexed and heated up to 98°C. The dissolved pellets were loaded onto TG gels. Electrophoresis was carried out in TG-SDS buffer for 1-1.5 hours at 100V. After electrophoresis, the proteins were transferred to nitrocellulose membranes using a commercial transfer kit. After transfer, the membranes were briefly rinsed in TBS, then blocked with 5% (w/v) milk powder in TBST for 45 minutes. The membranes were then rinsed in TBST and incubated with primary antibodies overnight at 4 °C with gentle rocking. Primary antibodies (Table S2) were diluted according to manufacturer instructions in TBST containing 5% BSA (w/v) and 0.04% sodium-azide (w/v). The following day the membranes were rinsed in TBST, then incubated with HRP-conjugated secondary antibodies diluted according to manufacturer instructions in a solution containing 5% milk powder in TBST. The membranes were then rinsed in TBST and dipped in an ECL mixture prior to exposure. Membranes were imaged using a Bio-Rad ChemiDoc MP imaging system.

#### Immunofluorescence imaging and analysis of cultured cells

Once ready for imaging, cells were fixed on the coverslips with 4% (v/v) formaldehyde solution in PBS for 20 minutes. The formaldehyde was then removed, and the coverslips were rinsed with PBS. Cells were then permeabilized with a 0.1% triton-X100 (v/v) solution in PBS for 20 minutes with gentle rocking. The triton solution was then removed, and the coverslips were rinsed with PBS. Next, coverslips were blocked with a 10% BSA (w/v) solution in PBS at 4°C overnight. The coverslips were then stained with primary antibodies (Table S2) diluted according to manufacturer instructions in a 1% BSA solution in PBS and incubated for 90 minutes at RT. The coverslips were then washed with PBS and incubated for 30 minutes with secondary antibodies, followed by washes with PBS and staining with DAPI working solution for 5 minutes. After DAPI staining the coverslips were washed with PBS and mounted on 25×75×1mm glass slides with ProLong Gold Antifade Mountant. The slides were visualized on a Nikon Eclipse Ti2 microscope with a Yokogawa CSU-W1 confocal scanner unit and photographed using a Photometrics Prime 95B CMOS camera. Nuclear area was measured manually using Fiji based on ImageJ version 1.53c available on imagej.net.

#### Mouse injection

SU-DIPG6 cells expressing GFP were stereotactically injected into the pons of male NGS mice as previously described (Venkatesh et al., 2019). Injection coordinates were: 0.8 mm posterior to lambda, 1 mm lateral to the sagittal suture, and 5 mm deep. Briefly, the skull of the mouse was exposed, and a small bore hole (0.5 mm) was made using a high-speed drill at the appropriate stereotactic coordinates. Approximately 300,000 cells in 3µl volume were injected at a speed of 0.3µL per minute into the pons with a 26-gauge Hamilton syringe; following 10 min pause, the needle was removed at a speed of 0.2 mm/minute. After closing the scalp, mice were placed on a warming pad and returned to their cages after full recovery. Mice were euthanized 14 weeks post-surgery and brains were removed for analysis.

#### Mouse slides collection and staining

Brains were collected, fixed in 4% paraformaldehyde, embedded in paraffin blocks in series of 6μm sections, with 50um skips made from the pons area. Slides were de-paraffinized and antigen retrieval was done using citric acid at pH6. Blocking for unspecific binding was done with 20% normal horse serum (NHS), 0.5% triton in PBS. Rabbit anti NUMA1 was diluted in 2% NHS and 0.5% triton and was incubated overnight. Slides were then incubated with an HRP conjugated goat anti rabbit 1:100 diluted 2% NHS for 1.5hr followed by 1:500 OPAL 690. Primary antibodies were stripped using microwave treatment with citric acid at pH6, blocked and again incubated with rabbit anti H3-K27M (1:100) and goat anti GFP (1:50). Donkey anti rabbit Cy3-conjugated and biotinylated were followed by a Sterptavidin-Cy2 incubation. Slides were imaged with Leica Mi8 microscope equipped with a motorized stage and a Leica DFC365 FX camera. Single x20 magnification images were tiled to receive a full scan of the tumor section. Slides were also imaged using a Nikon Eclipse Ti2 microscope with a Yokogawa CSU-W1 confocal scanner unit and photographed using a Photometrics Prime 95B CMOS camera.

#### Mouse slides image analysis

High magnification images were analyzed to detect H3-K27M signal heterogeneity. For each field of view, multiple optical Z sections were taken to cover the entire depth of the brain slice, and maximum projection images were produced. Nuclei were segmented using Fiji. Signal intensity histograms were generated by calculating the arcsinh transformed median intensity of the H3-K27M staining in each nucleus. Nuclei negative for NUMA staining were manually removed from the analysis to include only human cells.

#### Transfection with siRNA

Transfection with siRNA was done according to the manufacturer protocol. A day before transfection, 1.5M cells were seeded on a 10cm plate in DMEM without antibiotics. On the day of transfection, siRNA was diluted to a final concentration of 20nM in OptiMEM, and in a different tube 30μl of lipofectamine was mixed with 1470μl OptiMEM. Both mixtures were kept for 5 minutes at RT, and in the meantime the volume of medium in the 10cm plates was reduced to 7ml exactly. After 5 minutes of incubation at RT, the siRNA and lipofectamine in OptiMEM were mixed together and kept for 20 minutes at RT. A mix of 3ml was added to each 10cm plate, which was gently swirled to ensure consistent concentrations in the plate. The plates were incubated in a standard cell culture incubator at 37°C and 5% CO_2_ for 8 hours, and then the medium was changed to standard DMEM. The cells were harvested 48 hours after transfection.

### Quantification and statistical analysis

#### Nuclear size measurement

Group sizes are indicated in the figure legends. N was defined as the number of nuclei measured in each condition, and is stated in the figure legend. A one-sided Welch’s t-test was conducted to determine significance. The P-values *** p < 0.001 were considered statistically significant. Error bars represent the mean±SE.

#### CyTOF data quantification

The number of cells analyzed in each experiment is indicated in Table S3. N was defined as the number of cells in each condition. Welch’s t-test was conducted to determine significance when comparing differences in measured proteins, represented by box plots. A chi-square test was conducted when comparing fractions of cells in different conditions, represented by bar graphs. A confidence interval (CI) was inferred on correlations by bootstrapping, and Welch’s t-test was conducted to determine significance. For all tests in the paper, the P-value *** p < 0.001 was considered statistically significant.

## Notes

### Competing Interest Statement

The authors have declared no competing interest.

## References

Alabert, C., Barth, T.K., Reverón-Gómez, N., Sidoli, S., Schmidt, A., Jensen, O., Imhof, A., and Groth, A. (2015). Two distinct modes for propagation of histone PTMs across the cell cycle. Genes Dev. 29, 585–590.

Bartosovic, M., Kabbe, M., and Castelo-Branco, G. (2021). Single-cell CUT&Tag profiles histone modifications and transcription factors in complex tissues. Nat. Biotechnol.

Baslan, T., and Hicks, J. (2017). Unravelling biology and shifting paradigms in cancer with single-cell sequencing. Nat. Rev. Cancer 17, 557–569.

Bender, S., Tang, Y., Lindroth, A.M., Hovestadt, V., Jones, D.T.W., Kool, M., Zapatka, M., Northcott, P.A., Sturm, D., Wang, W., et al. (2013). Reduced H3K27me3 and DNA Hypomethylation Are Major Drivers of Gene Expression in K27M Mutant Pediatric High-Grade Gliomas. Cancer Cell 24, 660–672.

Boettiger, A.N., Bintu, B., Moffitt, J.R., Wang, S., Beliveau, B.J., Fudenberg, G., Imakaev, M., Mirny, L.A., Wu, C.T., and Zhuang, X. (2016). Super-resolution imaging reveals distinct chromatin folding for different epigenetic states. Nature 529, 418–422.

Brian, K., Ashot S. H., Shriya, D., and Nada, J. (2021). Polycomb repressive complex 2 in the driver’s seat of childhood and young adult brain tumours. Trends Cell Biol.

Brun, M., Jain, S., Monckton, E.A., and Godbout, R. (2018). Nuclear Factor I Represses the Notch Effector HEY1 in Glioblastoma. Neoplasia 20, 1023–1037.

Di Cerbo, V., Mohn, F., Ryan, D.P., Montellier, E., Kacem, S., Tropberger, P., Kallis, E., Holzner, M., Hoerner, L., Feldmann, A., et al. (2014). Acetylation of histone H3 at lysine 64 regulates nucleosome dynamics and facilitates transcription. Elife 2014, 1632.

Chan, K.M., Fang, D., Gan, H., Hashizume, R., Yu, C., Schroeder, M., Gupta, N., Mueller, S., David James, C., Jenkins, R., et al. (2013). The histone H3.3K27M mutation in pediatric glioma reprograms H3K27 methylation and gene expression. Genes Dev. 27, 985–990.

Chen, T., and Guestrin, C. (2016). XGBoost: A scalable tree boosting system. In Proceedings of the ACM SIGKDD International Conference on Knowledge Discovery and Data Mining, (New York, NY, USA: ACM), pp. 785–794.

Cheung, P., Vallania, F., Warsinske, H.C., Donato, M., Schaffert, S., Chang, S.E., Dvorak, M., Dekker, C.L., Davis, M.M., Utz, P.J., et al. (2018). Single-Cell Chromatin Modification Profiling Reveals Increased Epigenetic Variations with Aging. Cell 173, 1385–1397.e14.

Cusanovich, D.A., Daza, R., Adey, A., Pliner, H.A., Christiansen, L., Gunderson, K.L., Steemers, F.J., Trapnell, C., and Shendure, J. (2015). Multiplex single-cell profiling of chromatin accessibility by combinatorial cellular indexing. Science (80-.). 348, 910–914.

Dou, Y., Milne, T.A., Tackett, A.J., Smith, E.R., Fukuda, A., Wysocka, J., Allis, C.D., Chait, B.T., Hess, J.L., and Roeder, R.G. (2005). Physical association and coordinate function of the H3 K4 methyltransferase MLL1 and the H4 K16 acetyltransferase MOF. Cell 121, 873–885.

Duncan, E.M., Muratore-Schroeder, T.L., Cook, R.G., Garcia, B.A., Shabanowitz, J., Hunt, D.F., and Allis, C.D. (2008). Cathepsin L proteolytically processes histone H3 during mouse embryonic stem cell differentiation. Cell 135, 284–294.

Ernst, J., Kheradpour, P., Mikkelsen, T.S., Shoresh, N., Ward, L.D., Epstein, C.B., Zhang, X., Wang, L., Issner, R., Coyne, M., et al. (2011). Mapping and analysis of chromatin state dynamics in nine human cell types. Nature 473, 43–49.

Fedyuk, V., Erez, N., Furth, N., Beresh, O., Andreishcheva, E., Shinde, A., Jones, D., Zakai, B.B., Mavor, Y., Peretz, T., et al. (2021). Multiplexed Single-Molecule Epigenetic Analysis of Plasma-Isolated Nucleosomes for Cancer Diagnostics. BioRxiv.

Filbin, M., and Monje, M. (2019). Developmental origins and emerging therapeutic opportunities for childhood cancer. Nat. Med. 25, 367–376.

Filbin, M.G., Tirosh, I., Hovestadt, V., Shaw, M.L., Escalante, L.E., Mathewson, N.D., Neftel, C., Frank, N., Pelton, K., Hebert, C.M., et al. (2018). Developmental and oncogenic programs in H3K27M gliomas dissected by single-cell RNA-seq. Science (80-.). 360, 331–335.

Flavahan, W.A., Gaskell, E., and Bernstein, B.E. (2017). Epigenetic plasticity and the hallmarks of cancer. Science 357.

Furth, N., Algranati, D., Dassa, B., Beresh, O., Fedyuk, V., Morris, N., Kasper, L.H., Jones, D., Monje, M., Baker, S.J., et al. (2021). H3-K27M-Mutant Nucleosomes Interact with MLL1 to Shape the Glioma Epigenetic Landscape. BioRxiv.

Grasso, C.S., Tang, Y., Truffaux, N., Berlow, N.E., Liu, L., Debily, M.A., Quist, M.J., Davis, L.E., Huang, E.C., Woo, P.J., et al. (2015). Functionally defined therapeutic targets in diffuse intrinsic pontine glioma. Nat. Med. 21, 555–559.

Grosselin, K., Durand, A., Marsolier, J., Poitou, A., Marangoni, E., Nemati, F., Dahmani, A., Lameiras, S., Reyal, F., Frenoy, O., et al. (2019). High-throughput single-cell ChIP-seq identifies heterogeneity of chromatin states in breast cancer. Nat. Genet. 51, 1060–1066.

Han, G., Spitzer, M.H., Bendall, S.C., Fantl, W.J., and Nolan, G.P. (2018). Metal-isotope-tagged monoclonal antibodies for high-dimensional mass cytometry. Nat. Protoc. 2018 1310 13, 2121–2148.

Harutyunyan, A.S., Chen, H., Lu, T., Horth, C., Nikbakht, H., Krug, B., Russo, C., Bareke, E., Marchione, D.M., Coradin, M., et al. (2020). H3K27M in Gliomas Causes a One-Step Decrease in H3K27 Methylation and Reduced Spreading within the Constraints of H3K36 Methylation. Cell Rep. 33.

Ichijima, Y., Sakasai, R., Okita, N., Asahina, K., Mizutani, S., and Teraoka, H. (2005). Phosphorylation of histone H2AX at M phase in human cells without DNA damage response. Biochem. Biophys. Res. Commun. 336, 807–812.

Jaitin, D.A., Kenigsberg, E., Keren-Shaul, H., Elefant, N., Paul, F., Zaretsky, I., Mildner, A., Cohen, N., Jung, S., Tanay, A., et al. (2014). Massively parallel single-cell RNA-seq for marker-free decomposition of tissues into cell types. Science 343, 776–779.

Jeon, H.M., Jin, X., Lee, J.S., Oh, S.Y., Sohn, Y.W., Park, H.J., Kyeung, M.J., Park, W.Y., Nam, D.H., DePinho, R.A., et al. (2008). Inhibitor of differentiation 4 drives brain tumor-initiating cell genesis through cyclin E and notch signaling. Genes Dev. 22, 2028–2033.

Kelsey, G., Stegle, O., and Reik, W. (2017). Single-cell epigenomics: Recording the past and predicting the future. Science 358, 69–75.

Kheir, T.B., and Lund, A.H. (2010). Epigenetic dynamics across the cell cycle. Essays Biochem. 48, 107–120.

Kim, K., Punj, V., Kim, J.-M., Lee, S., Ulmer, T.S., Lu, W., Rice, J.C., and An, W. (2016). MMP-9 facilitates selective proteolysis of the histone H3 tail at genes necessary for proficient osteoclastogenesis. Genes Dev. 30, 208–219.

Klein, K.N., Zhao, P.A., Lyu, X., Sasaki, T., Bartlett, D.A., Singh, A.M., Tasan, I., Zhang, M., Watts, L.P., Hiraga, S.I., et al. (2021). Replication timing maintains the global epigenetic state in human cells. Science (80-.). 372, 371–378.

Krug, B., De Jay, N., Harutyunyan, A.S., Deshmukh, S., Marchione, D.M., Guilhamon, P., Bertrand, K.C., Mikael, L.G., McConechy, M.K., Chen, C.C.L., et al. (2019). Pervasive H3K27 Acetylation Leads to ERV Expression and a Therapeutic Vulnerability in H3K27M Gliomas. Cancer Cell 35, 782–797.e8.

Lewis, P.W., Müller, M.M., Koletsky, M.S., Cordero, F., Lin, S., Banaszynski, L.A., Garcia, B.A., Muir, T.W., Becher, O.J., and Allis, C.D. (2013). Inhibition of PRC2 activity by a gain-of-function H3 mutation found in pediatric glioblastoma. Science (80-.). 340, 857–861.

Luo, C., Hajkova, P., and Ecker, J.R. (2018). Dynamic DNA methylation: In the right place at the right time. Science 361, 1336–1340.

McInnes, L., Healy, J., Saul, N., and Großberger, L. (2018). UMAP: Uniform Manifold Approximation and Projection. J. Open Source Softw. 3, 861.

Nacev, B.A., Feng, L., Bagert, J.D., Lemiesz, A.E., Gao, J., Soshnev, A.A., Kundra, R., Schultz, N., Muir, T.W., and Allis, C.D. (2019). The expanding landscape of “oncohistone” mutations in human cancers. Nature 567, 473–478.

Neftel, C., Laffy, J., Filbin, M.G., Hara, T., Shore, M.E., Rahme, G.J., Richman, A.R., Silverbush, D., Shaw, M.L., Hebert, C.M., et al. (2019). An Integrative Model of Cellular States, Plasticity, and Genetics for Glioblastoma. Cell 178, 835–849.e21.

Pajovic, S., Siddaway, R., Bridge, T., Sheth, J., Rakopoulos, P., Kim, B., Ryall, S., Agnihotri, S., Phillips, L., Yu, M., et al. (2020). Epigenetic activation of a RAS/MYC axis in H3.3K27M-driven cancer. Nat. Commun. 11.

Palii, C.G., Cheng, Q., Gillespie, M.A., Shannon, P., Mazurczyk, M., Napolitani, G., Price, N.D., Ranish, J.A., Morrissey, E., Higgs, D.R., et al. (2019). Single-Cell Proteomics Reveal that Quantitative Changes in Co-expressed Lineage-Specific Transcription Factors Determine Cell Fate. Cell Stem Cell 24, 812.

Phillips, R.E., Soshnev, A.A., and Allis, C.D. (2020). Epigenomic Reprogramming as a Driver of Malignant Glioma. Cancer Cell 38, 647–660.

Piunti, A., Hashizume, R., Morgan, M.A., Bartom, E.T., Horbinski, C.M., Marshall, S.A., Rendleman, E.J., Ma, Q., Takahashi, Y.H., Woodfin, A.R., et al. (2017). Therapeutic targeting of polycomb and BET bromodomain proteins in diffuse intrinsic pontine gliomas. Nat. Med. 23, 493–500.

Reverón-Gómez, N., González-Aguilera, C., Stewart-Morgan, K.R., Petryk, N., Flury, V., Graziano, S., Johansen, J.V., Jakobsen, J.S., Alabert, C., and Groth, A. (2018). Accurate Recycling of Parental Histones Reproduces the Histone Modification Landscape during DNA Replication. Mol. Cell 72, 239–249.e5.

Rice, J.C., and Allis, C.D. (2001). Histone methylation versus histone acetylation: New insights into epigenetic regulation. Curr. Opin. Cell Biol. 13, 263–273.

Rotem, A., Ram, O., Shoresh, N., Sperling, R.A., Goren, A., Weitz, D.A., and Bernstein, B.E. (2015). Single-cell ChIP-seq reveals cell subpopulations defined by chromatin state. Nat. Biotechnol. 33, 1165–1172.

Schachner, L.F., Jooß, K., Morgan, M.A., Piunti, A., Meiners, M.J., Kafader, J.O., Lee, A.S., Iwanaszko, M., Cheek, M.A., Burg, J.M., et al. (2021). Decoding the protein composition of whole nucleosomes with Nuc-MS. Nat. Methods 18, 303–308.

Schwartzentruber, J., Korshunov, A., Liu, X.Y., Jones, D.T.W., Pfaff, E., Jacob, K., Sturm, D., Fontebasso, A.M., Quang, D.A.K., Tönjes, M., et al. (2012). Driver mutations in histone H3.3 and chromatin remodelling genes in paediatric glioblastoma. Nature 482, 226–231.

Sharko, J., Grinstein, G., and Marx, K.A. (2008). Vectorized radviz and its application to multiple cluster datasets. In IEEE Transactions on Visualization and Computer Graphics, pp. 1444–1451.

Shema, E., Jones, D., Shoresh, N., Donohue, L., Ram, O., and Bernstein, B.E. (2016). Single-molecule decoding of combinatorially modified nucleosomes. Science (80-.). 352, 717–721.

Shema, E., Bernstein, B.E., and Buenrostro, J.D. (2019). Single-cell and single-molecule epigenomics to uncover genome regulation at unprecedented resolution. Nat. Genet. 51, 19–25.

Shirane, K., Miura, F., Ito, T., and Lorincz, M.C. (2020). NSD1-deposited H3K36me2 directs de novo methylation in the mouse male germline and counteracts Polycomb-associated silencing. Nat. Genet. 2020 5210 52, 1088–1098.

Silveira, A.B., Kasper, L.H., Fan, Y., Jin, H., Wu, G., Shaw, T.I., Zhu, X., Larson, J.D., Easton, J., Shao, Y., et al. (2019). H3.3 K27M Depletion Increases Differentiation and Extends Latency of Diffuse Intrinsic Pontine Glioma Growth In Vivo. Acta Neuropathol. 137, 637.

Smallwood, S.A., Lee, H.J., Angermueller, C., Krueger, F., Saadeh, H., Peat, J., Andrews, S.R., Stegle, O., Reik, W., and Kelsey, G. (2014). Single-cell genome-wide bisulfite sequencing for assessing epigenetic heterogeneity. Nat. Methods 11, 817–820.

Smith, E.R., Cayrou, C., Huang, R., Lane, W.S., Côté, J., and Lucchesi, J.C. (2006). A Human Protein Complex Homologous to the Drosophila MSL Complex Is Responsible for the Majority of Histone H4 Acetylation at Lysine 16. Mol. Cell. Biol. 26, 387–387.

Suvà, M.L., and Tirosh, I. (2019). Single-Cell RNA Sequencing in Cancer: Lessons Learned and Emerging Challenges. Mol. Cell 75, 7–12.

Suvà, M.L., Rheinbay, E., Gillespie, S.M., Patel, A.P., Wakimoto, H., Rabkin, S.D., Riggi, N., Chi, A.S., Cahill, D.P., Nahed, B. V., et al. (2014). Reconstructing and reprogramming the tumor-propagating potential of glioblastoma stem-like cells. Cell 157, 580–594.

Tiscornia, G., Singer, O., and Verma, I.M. (2006). Production and purification of lentiviral vectors. Nat. Protoc. 2006 11 1, 241–245.

Torres, C.M., Biran, A., Burney, M.J., Patel, H., Henser-Brownhill, T., Cohen, A.H.S., Li, Y., Ben-Hamo, R., Nye, E., Spencer-Dene, B., et al. (2016). The linker histone H1.0 generates epigenetic and functional intratumor heterogeneity. Science (80-.). 353.

Valencia, A.M., and Kadoch, C. (2019). Chromatin regulatory mechanisms and therapeutic opportunities in cancer. Nat. Cell Biol. 21, 152–161.

Venkatesh, H.S., Morishita, W., Geraghty, A.C., Silverbush, D., Gillespie, S.M., Arzt, M., Tam, L.T., Espenel, C., Ponnuswami, A., Ni, L., et al. (2019). Electrical and synaptic integration of glioma into neural circuits. Nature 573, 539–545.

Wallace, G.C., Dixon-Mah, Y.N., Vandergrift, W.A., Ray, S.K., Haar, C.P., Mittendorf, A.M., Patel, S.J., Banik, N.L., Giglio, P., and Das, A. (2013). Targeting Oncogenic ALK and MET: A Promising Therapeutic Strategy for Glioblastoma. Metab. Brain Dis. 28, 355.

Wang, X., Prager, B.C., Wu, Q., Kim, L.J.Y., Gimple, R.C., Shi, Y., Yang, K., Morton, A.R., Zhou, W., Zhu, Z., et al. (2018). Reciprocal Signaling between Glioblastoma Stem Cells and Differentiated Tumor Cells Promotes Malignant Progression. Cell Stem Cell 22, 514–528.e5.

Weinberg, D.N., Papillon-Cavanagh, S., Chen, H., Yue, Y., Chen, X., Rajagopalan, K.N., Horth, C., McGuire, J.T., Xu, X., Nikbakht, H., et al. (2019). The histone mark H3K36me2 recruits DNMT3A and shapes the intergenic DNA methylation landscape. Nat. 2019 5737773 573, 281–286.

Wimmers, F., Donato, M., Kuo, A., Ashuach, T., Gupta, S., Li, C., Dvorak, M., Foecke, M.H., Chang, S.E., De Jong, S.E., et al. (2021). Single-cell analysis of the epigenomic and transcriptional landscape of innate immunity to seasonal and adjuvanted pandemic influenza vaccination in humans. MedRxiv 2021.05.24.21253087.

Woodworth, M.A., Ng, K.K.H., Halpern, A.R., Pease, N.A., Nguyen, P.H.B., Kueh, H.Y., and Vaughan, J.C. (2021). Multiplexed single-cell profiling of chromatin states at genomic loci by expansion microscopy Marcus. Nucleic Acids Res. 1–15.

Wu, G., Broniscer, A., McEachron, T.A., Lu, C., Paugh, B.S., Becksfort, J., Qu, C., Ding, L., Huether, R., Parker, M., et al. (2012). Somatic histone H3 alterations in pediatric diffuse intrinsic pontine gliomas and non-brainstem glioblastomas. Nat. Genet. 44, 251–253.

Wu, S.J., Furlan, S.N., Mihalas, A.B., Kaya-Okur, H.S., Feroze, A.H., Emerson, S.N., Zheng, Y., Carson, K., Cimino, P.J., Keene, C.D., et al. (2021). Single-cell CUT&Tag analysis of chromatin modifications in differentiation and tumor progression. Nat. Biotechnol.

Wu, X., Wakefield, J.K., Liu, H., Xiao, H., Kralovics, R., Prchal, J.T., and Kappes, J.C. (2000). Development of a novel trans-lentiviral vector that affords predictable safety. Mol. Ther. 2, 47–55.

Xu, J., Ma, H., Jin, J., Uttam, S., Fu, R., Huang, Y., and Liu, Y. (2018). Super-Resolution Imaging of Higher-Order Chromatin Structures at Different Epigenomic States in Single Mammalian Cells. Cell Rep. 24, 873–882.

Ying, M., Tilghman, J., Wei, Y., Guerrero-Cazares, H., Quinones-Hinojosa, A., Ji, H., and Laterra, J. (2014). Kruppel-like Factor-9 (KLF9) Inhibits Glioblastoma Stemness through Global Transcription Repression and Integrin α6 Inhibition. J. Biol. Chem. 289, 32742.

Zhang, X., and Zhang, Z. (2019). Oncohistone Mutations in Diffuse Intrinsic Pontine Glioma. Trends in Cancer 5, 799–808.

Zhang, B., Xu, C., Liu, J., Yang, J., Gao, Q., and Ye, F. (2021). Nidogen-1 expression is associated with overall survival and temozolomide sensitivity in low-grade glioma patients. Aging (Albany. NY). 13, 9085–9107.

Zhang, Y., Jang, Y., Lee, J.E., Ahn, J.W., Xu, L., Holden, M.R., Cornett, E.M., Krajewski, K., Klein, B.J., Wang, S.P., et al. (2019). Selective binding of the PHD6 finger of MLL4 to histone H4K16ac links MLL4 and MOF. Nat. Commun. 10.

Zhao, Z., and Shilatifard, A. (2019). Epigenetic modifications of histones in cancer. Genome Biol. 20.

Zhou, P., Wu, E., Alam, H.B., and Li, Y. (2014). Histone Cleavage as a Mechanism for Epigenetic Regulation: Current Insights and Perspectives. Curr. Mol. Med. 14, 1164.

Zhou, W., Jiang, D., Tian, J., Liu, L., Lu, T., Huang, X., and Sun, H. (2019). Acetylation of H3K4, H3K9, and H3K27 mediated by p300 regulates the expression of GATA4 in cardiocytes. Genes Dis. 6, 318–325.

