## Supplementary information for "Single-cell epigenetic analysis reveals principles of chromatin states in H3.3-K27M gliomas"

Fig. S1

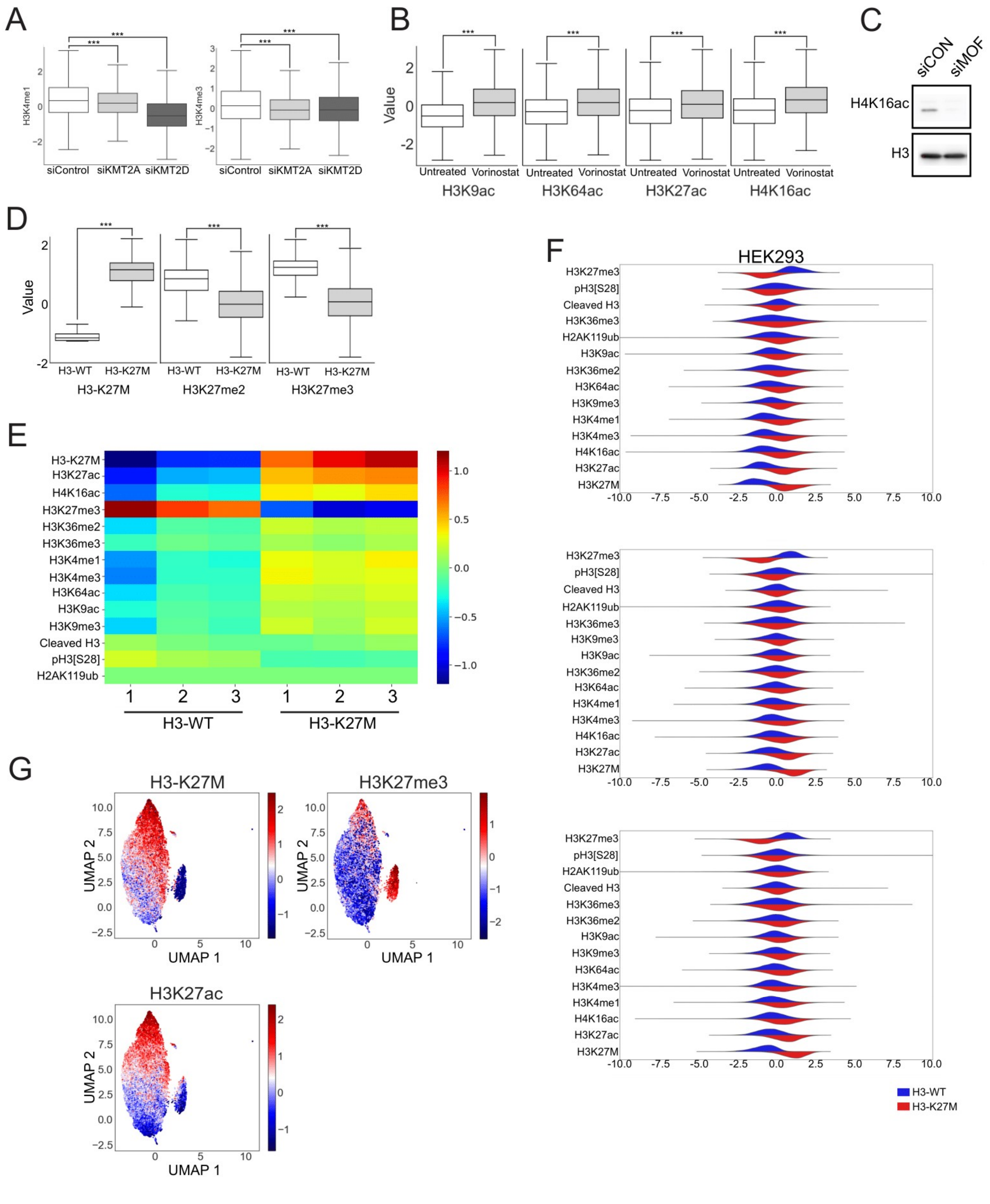
A
H3K4me1
siControl
siKMT2A
siKMT2D
H3K4me3
B
Value
Untreated
Vorinostat
H3K9ac
H3K64ac
H3K27ac
H4K16ac
C
siCON
siMOF
H4K16ac
H3
D
H3-WT
H3-K27M
H3-K27M
H3K27me2
H3K27me3
E
H3-K27M
H3K27ac
H4K16ac
H3K27me3
H3K36me2
H3K36me3
H3K4me1
H3K4me3
H3K64ac
H3K9ac
H3K9me3
Cleaved H3
pH3[S28]
H2AK119ub
H3-WT
H3-K27M
F
HEK293
H3-WT
H3-K27M
G
H3-K27M
H3K27me3
H3K27ac
UMAP 1
UMAP 2

**Figure S1: High reproducibility of H3-K27M-mediated epigenetic alterations in HEK293 cells. Related to Figure 1.**

(**A-D**) Validations of antibodies included in the CyTOF panel, by CyTOF (A,B,D) or by Western blot (C). (**A**) Expression levels of H3K4me1 (left) and H3K4me3 (right) in HEK293 cells following siRNA-mediated knock-down of KMT2A or KMT2D, the enzymes that deposit these marks. P values were calculated by Welch's t-test. *** P value < 0.001. (**B**) Expression levels of histone acetylations in SU-DIPG13 treated with the histone deacetylase inhibitor Vorinostat for 96 hours, compared to untreated cells. Significant elevation is detected for all acetylations, as expected. P values were calculated as in (A). **(C)** Western blot for H4K16ac in HEK293 cells following siRNA-mediated knock-down of the H4K16ac acetyltransferase, MOF. H3 is used as loading control. **(D)** Expression levels of H3K27M, H3K27me2 and H3K27me3 in WT DIPG cells (SJ-HGGX39) ectopically expressing WT H3.3 or H3.3-K27M. P values were calculated as in (A). (**E**) Heat map matrix of the expression levels of the indicated histone modifications in HEK293 cells induced to express H3-WT or H3-K27M. Numbers represent three independent biological repeats. Induction period was 10, 7 and 4 days in experiments 1, 2 and 3, respectively. Epigenetic alterations are highly reproducible despite the different induction times. (**F**) Data distribution of the indicated epigenetic modifications in HEK293 cells expressing WT H3.3 or H3.3-K27M. Top: 10 days of H3-K27M induction. Middle: 7 days induction. Bottom: 4 days induction. (**G**) UMAP analysis of the HEK293 cells induced to express H3-K27M. Shown is the small cluster of cells that failed to express H3-K27M, as indicated by the lack of H3-K27M signal as well as lack of effect on H3K27me3 and H3K27ac. The exact panel composition used for each experiment, and the number of cells analyzed, is described in Table S2 and Table S3, respectively.

Fig. S2

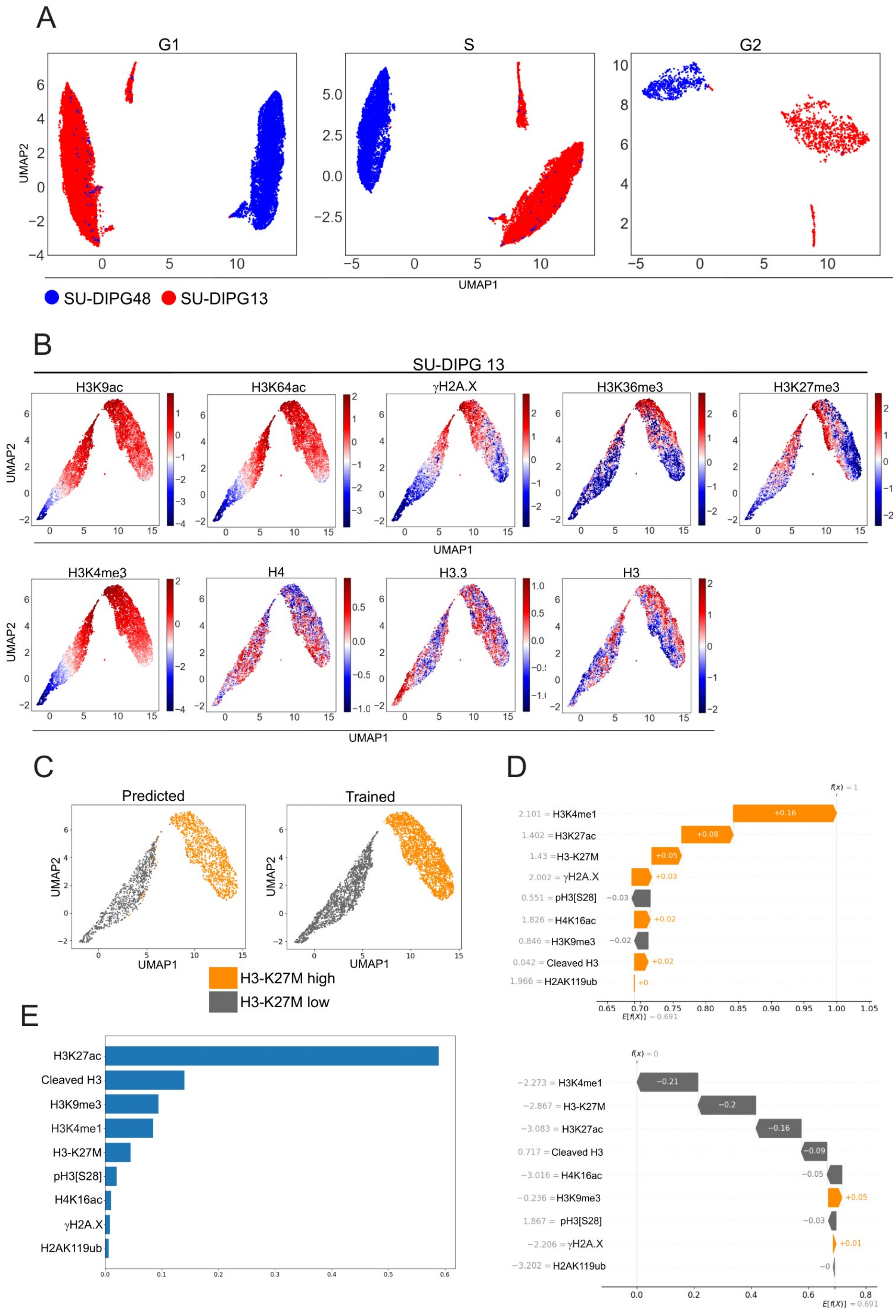
A
G1
S
G2
UMAP2
UMAP1
SU-DIPG48
SU-DIPG13
B
SU-DIPG 13
H3K9ac
H3K64ac
γH2A.X
H3K36me3
H3K27me3
H3K4me3
H4
H3.3
H3
UMAP2
UMAP1
C
Predicted
Trained
UMAP2
UMAP1
H3-K27M high
H3-K27M low
D
f(x) = 1
2.101 = H3K4me1
1.402 = H3K27ac
1.43 = H3-K27M
2.002 = γH2A.X
0.551 = pH3[S28]
1.826 = H4K16ac
0.846 = H3K9me3
0.042 = Cleaved H3
1.966 = H2AK119ub
+0.16
+0.08
+0.05
+0.03
−0.03
+0.02
−0.02
+0.02
+0
E[f(X)] = 0.691
f(x) = 0
−2.273 = H3K4me1
−2.867 = H3-K27M
−3.083 = H3K27ac
0.717 = Cleaved H3
−3.016 = H4K16ac
−0.236 = H3K9me3
1.867 = pH3[S28]
−2.206 = γH2A.X
−3.202 = H2AK119ub
−0.21
−0.2
−0.16
−0.09
−0.05
+0.05
−0.03
+0.01
−0
E[f(X)] = 0.691
E
H3K27ac
Cleaved H3
H3K9me3
H3K4me1
H3-K27M
pH3[S28]
H4K16ac
γH2A.X
H2AK119ub

Fig. S2 continued

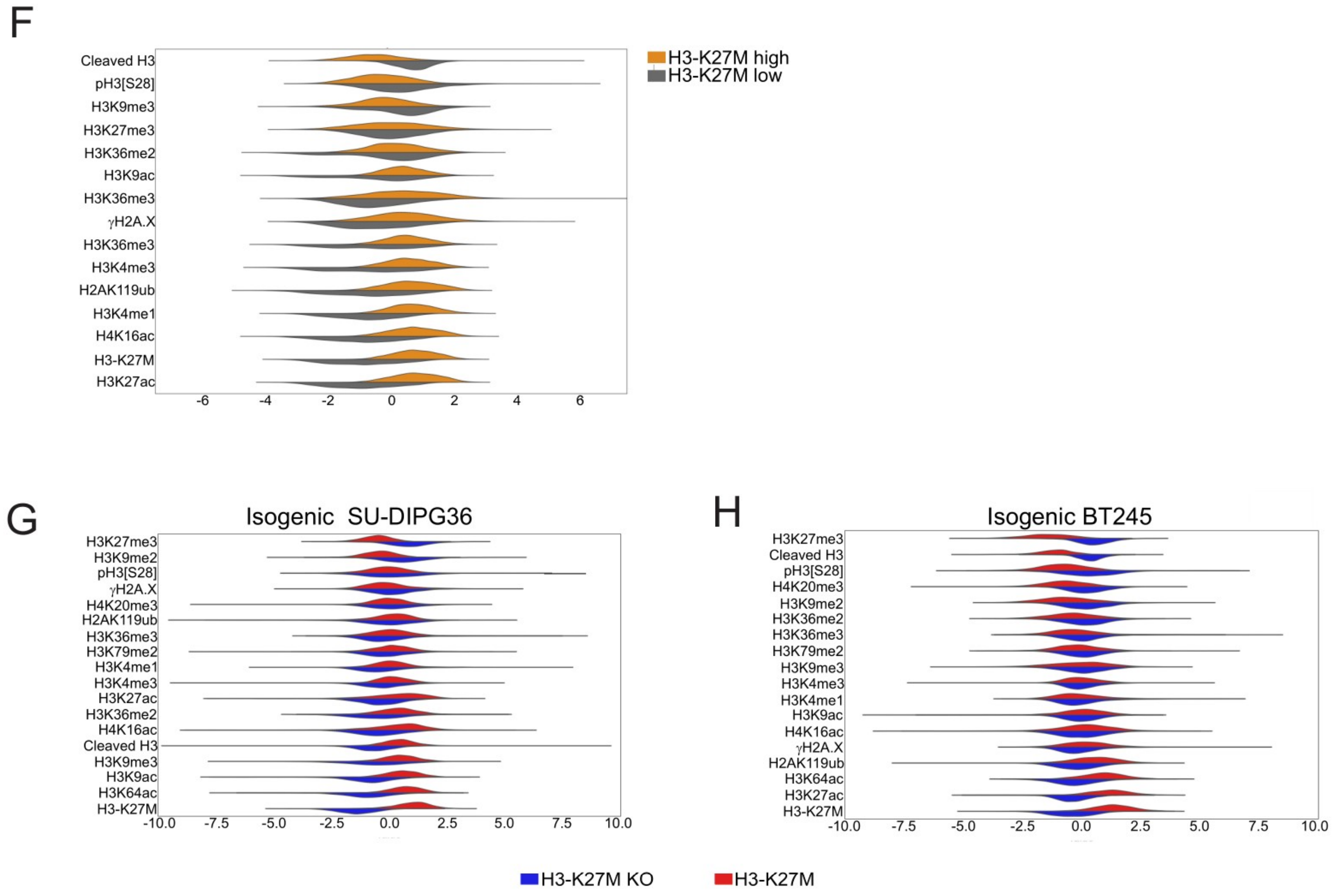

**Figure S2: Machine learning to identify epigenetic modifications influencing the two epigenetic states in SU-DIPG13. Related to Figure 2.**

(**A**) SU-DIPG48 tumor-derived cells expressing WT H3 (blue) and SU-DIPG13 expressing H3-K27M (red) were analyzed by CyTOF. Cell cycle markers, included in the CyTOF panel, were used to determine the cell cycle phase of each cell: G1, S and G2. For each phase, a joint UMAP of the two cell lines was generated, based on all epigenetic modifications. The two tumor-derived lines cluster separately in each phase of the cell cycle (red/blue indicates the sample's index/barcode). A single repeat was done for this experiment. For panel composition and number of cells analyzed, see Tables S2-3. (**B**) UMAP of SU-DIPG13 cells based on epigenetic modifications as in figure 3B, showing two distinct clusters. Levels of the indicated modifications in these two clusters are shown. **(C-E**) Machine learning applied to study the allocation of cells to the two epigenetic clusters in SU-DIPG13, using gradient boosting algorithm. (**C**) UMAP representation of the trained and predicted datasets. Orange and grey relate to the H3-K27M-high and H3-K27M-low clusters, respectively. (**D**) Two representative waterfall plots; for each instance the figure shows the Shapley value[54] of each of the modifications. Modifications shifting the result toward the H3-K27M-low cluster are denoted in gray, while those shifting in the direction of the H3-K27M-high cluster are denoted in orange. The modifications are ordered by decreasing Shapley values, where the most contributing modifications (largest absolute value) are shown at the top of each plot. (**E**) The mean absolute Shapley values for the various modifications over all instances in the training set. The value shown indicates the importance of the modification to the classification. (**F**) Data distribution of the epigenetic modifications, as measured by CyTOF, for the two epigenetic clusters identified in SU-DIPG13. (**G-H**) Data distribution of the epigenetic modifications, as measured by CyTOF, in SU-DIPG36 (G) and BT245 (H) cell lines, in the control cells expressing H3K27M versus cells knocked-out (KO) for the mutant histone. For panel composition and number of cells analyzed, see Tables S2-3. A single repeat was done per cell line.

Fig. S3

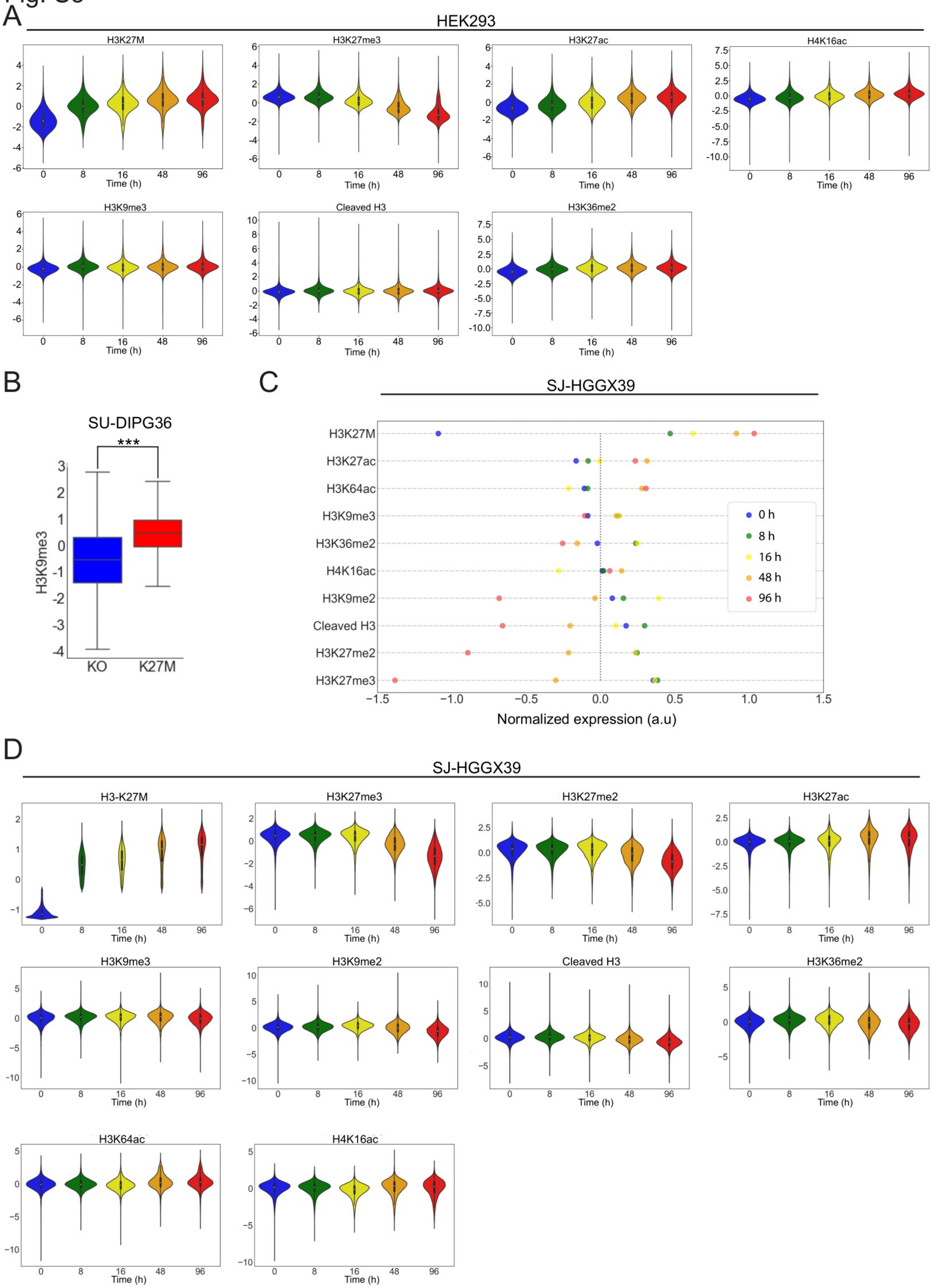
A
HEK293
H3K27M
H3K27me3
H3K27ac
H4K16ac
H3K9me3
Cleaved H3
H3K36me2
Time (h)
0
8
16
48
96
B
SU-DIPG36
***
H3K9me3
KO
K27M
C
SJ-HGGX39
H3K27M
H3K27ac
H3K64ac
H3K9me3
H3K36me2
H4K16ac
H3K9me2
Cleaved H3
H3K27me2
H3K27me3
0 h
8 h
16 h
48 h
96 h
Normalized expression (a.u)
D
SJ-HGGX39
H3-K27M
H3K27me3
H3K27me2
H3K27ac
H3K9me3
H3K9me2
Cleaved H3
H3K36me2
H3K64ac
H4K16ac
Time (h)

**Figure S3: Dynamics of H3-K27M-induced epigenetic alterations. Related to Figure 3.**

**(A)** HEK293 cells were induced to express H3-K27M for 8, 16, 48 and 96 hours, compared to cells expressing WT-H3 (marked as zero induction of H3-K27M), and analyzed by CyTOF . Violin plots of the indicated modifications show alterations in their distribution following H3K27M induction. To calculate the significance of changes over time, we performed Welch's t-test for each modification at each of the time points, relative to the time point that preceded it. All changes are significant (p-value < 0.001) except for H3K27me3 levels at time points 0 and 8, which remain the same. The experiment was performed once with multiple time points. **(B)** H3K9me3 expression levels in SU-DIPG36 cell line, as measured by CyTOF, in the control cells expressing H3K27M versus cells knocked-out (KO) for the mutant histone. P values were calculated by Welch's t-test. *** P value < 0.001. (**C-D**) SJ-HGGX39 cells were induced to express H3-K27M for 8, 16, 48 and 96 hours, compared to non-induced cells. CyTOF analysis was done as in A, with the addition of antibodies targeting H3K27me2 and H3K9me2. Shown are the mean values (C) and violin plots (D) of the indicated modifications at different time points following H3-K27M induction. P values were calculated as in (A): all changes are significant except for the change in H3K27me2 levels from 0 to 8, and 8 to 16 hours; the H3K36me2 change from 8 to 16 hours; the H4K16ac change from 0 to 8 hours, and the H3K9me3 change from 8 to 16 hours, and 16 to 48 hours. The experiment was performed once with multiple time points. For panel composition and number of cells analyzed, see Tables S2-3.

Fig. S4

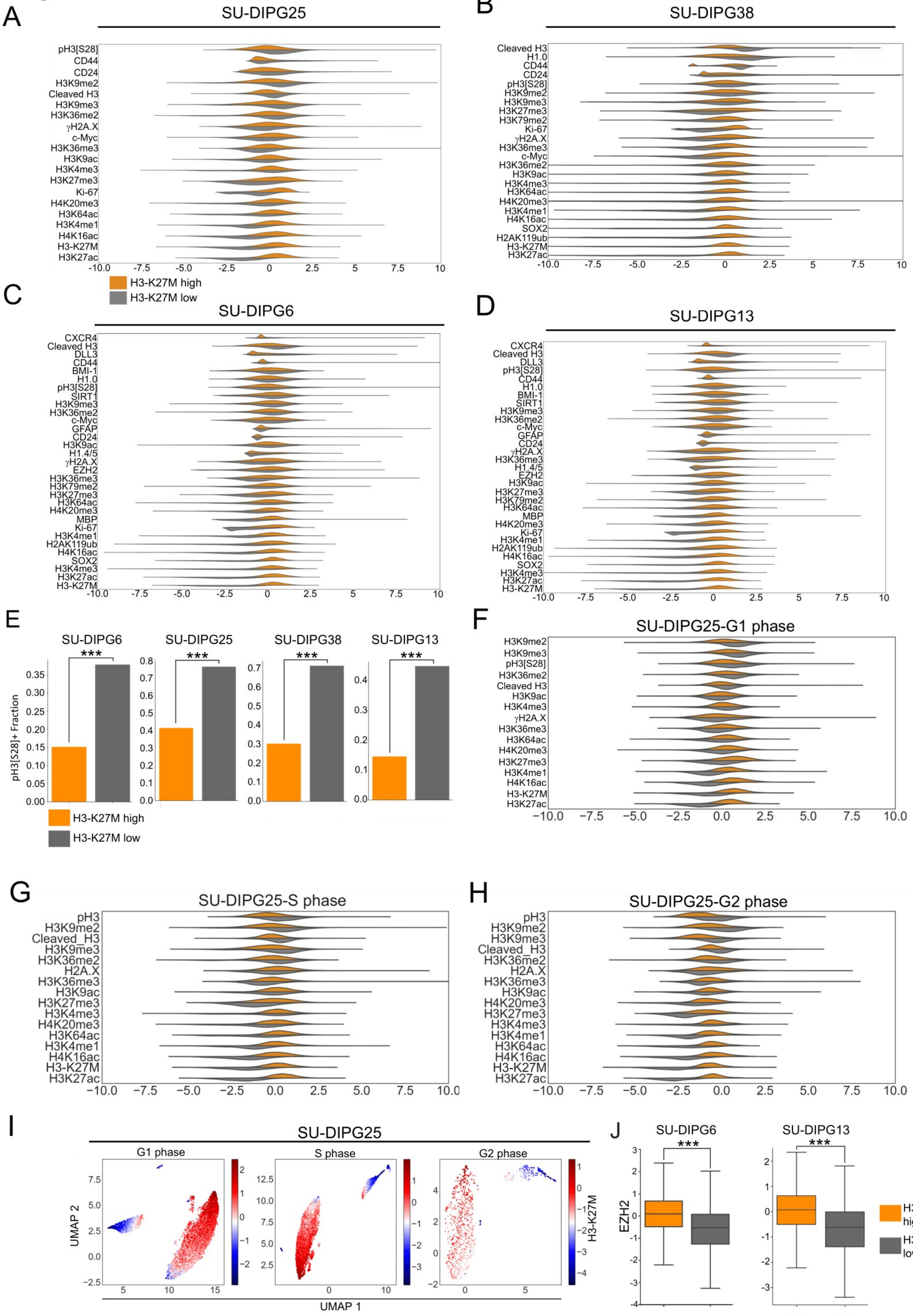
A
SU-DIPG25
pH3[S28]
CD44
CD24
H3K9me2
Cleaved H3
H3K9me3
H3K36me2
γH2A.X
c-Myc
H3K36me3
H3K9ac
H3K4me3
H3K27me3
Ki-67
H4K20me3
H3K64ac
H3K4me1
H4K16ac
H3-K27M
H3K27ac
H3-K27M high
H3-K27M low
B
SU-DIPG38
Cleaved H3
H1.0
CD44
CD24
pH3[S28]
H3K9me2
H3K9me3
H3K27me3
H3K79me2
Ki-67
γH2A.X
H3K36me3
c-Myc
H3K36me2
H3K9ac
H3K4me3
H3K64ac
H4K20me3
H3K4me1
H4K16ac
SOX2
H2AK119ub
H3-K27M
H3K27ac
C
SU-DIPG6
D
SU-DIPG13
E
SU-DIPG6
SU-DIPG25
SU-DIPG38
SU-DIPG13
pH3[S28]+ Fraction
***
H3-K27M high
H3-K27M low
F
SU-DIPG25-G1 phase
G
SU-DIPG25-S phase
H
SU-DIPG25-G2 phase
I
SU-DIPG25
G1 phase
S phase
G2 phase
UMAP 2
UMAP 1
H3-K27M
J
SU-DIPG6
SU-DIPG13
EZH2
***
H3-K27M high
H3-K27M low

Fig. S4 continued

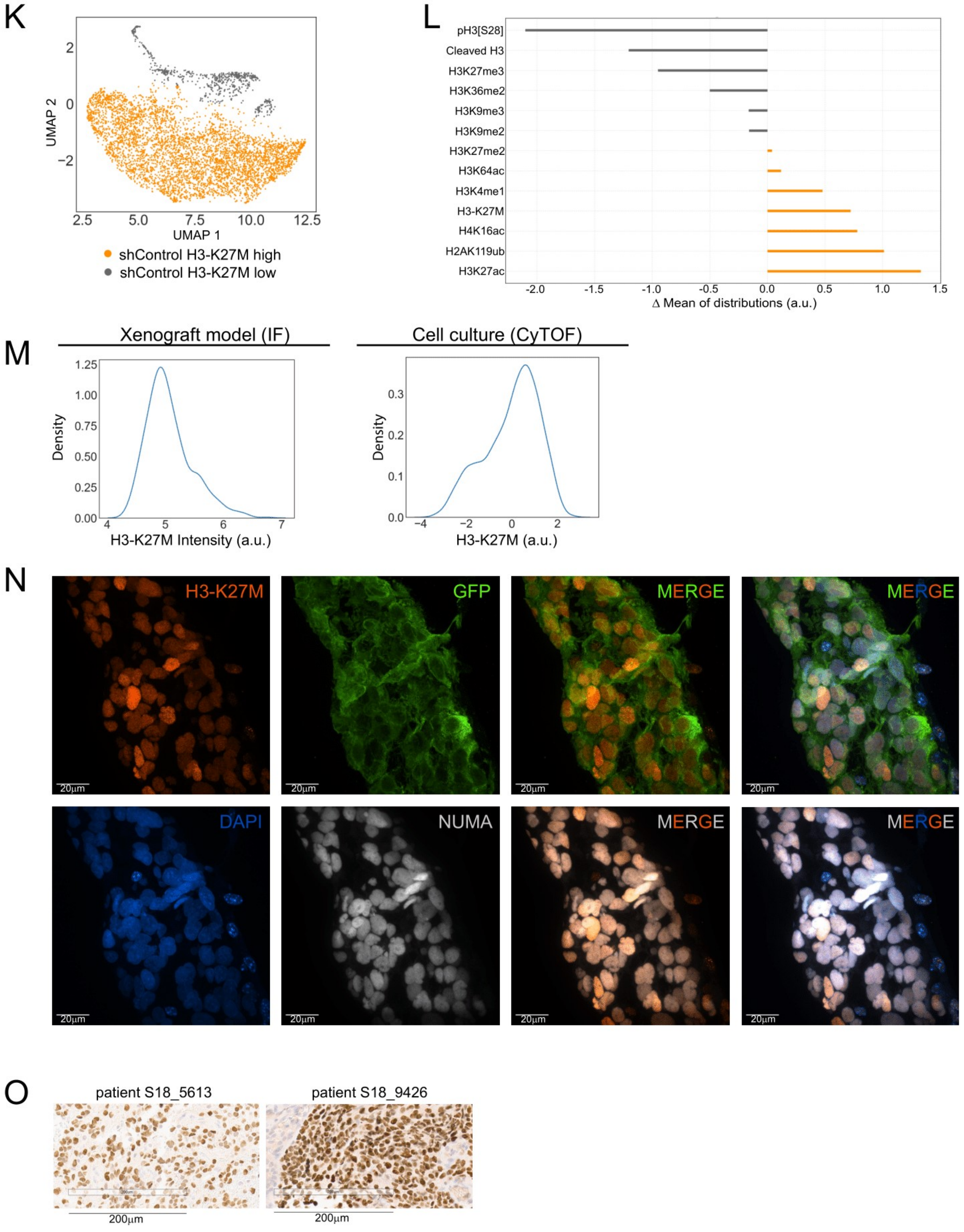

**Figure S4: The two epigenetic states are robust across several patient-derived lines, with H3-K27M heterogeneity observed *in-vivo* in human samples. Related to Figure 4.**

(**A-D**) Data distribution of the epigenetic modifications, as measured by CyTOF, for the two epigenetic clusters identified in **(A)** SU-DIPG 25, **(B)** SU-DIPG38, **(C)** SU-DIPG6 and **(D)** SU-DIPG13. Presented is one biological representative repeat out of four CyTOF experiments for SU-DIPG13, three for SU-DIPG25, one for SU-DIPG6 and one for SU-DIPG38. (**E**) Fraction of cells positive for the mitosis-associated phosphorylation pH3[S28] in the two epigenetic clusters, in the indicated DIPG lines. P values were calculated by chi square test. *** P value < 0.001. (**F-I**) SU-DIPG25 tumor-derived cells were analyzed by CyTOF. Cell cycle markers, included in the CyTOF panel, were used to determine the cell cycle phase of each cell: G1, S and G2. (**F-H**) Data distribution of the epigenetic modifications, as measured by CyTOF, for the two epigenetic clusters, for cells in **(F)** G1 phase, **(G)** S phase and **(H)** G2 phase. **(I)** UMAP analysis, based on all epigenetic modifications, indicates that the two epigenetic subpopulations are distinct at all phases of the cell cycle. Shown are normalized levels of H3K27M. (**J**) EZH2 expression levels as measured by CyTOF in the two epi-clusters of SU-DIPG6 and SU-DIPG13. P values were calculated by Welch's t-test. *** P value < 0.001. (**K**) SU-DIPG13 cells were infected with shRNA targeting the H3F3B WT gene of H3.3 ('shControl', part of the experiment presented in Figure 4H), and analyzed by CyTOF. UMAP was performed based on all epigenetic markers. Gray and orange colors indicate the H3-K27M-low and H3-K27M-high clusters, respectively. (**L**) Fold change differences between the two clusters identified in (K) for the indicated epigenetic modifications. Mean values (after transformation, scaling and normalization) of the H3-K27M-low cluster were subtracted from the H3-K27M-high cluster. (**M**) Left: density plot of the arcsinh transformed median values of nuclear H3-K27M intensity, measured by immunohistochemistry for H3-K27M, in DIPG tumors *in-vivo* in the mouse xenograft model. Shown is one representative repeat out of three mice that were analyzed. Right: density plot of H3-K27M levels (after transformation, scaling and normalization), as measured by CyTOF in SU-DIPG13 cells. (**N**) Confocal images of DIPG tumors *in-vivo* in the mouse xenograft model, stained for DAPI (blue), GFP (green), H3-K27M (red) and NUMA (white). Co-staining indicates that H3K27M positive cells are also positive for human specific NUMA and GFP staining. Scale bar is 20µm. (**O**) Human patients' autopsy samples, stained for H3-K27M (similar to Figure 4J). Scale bar = 200µm.

Fig. S5

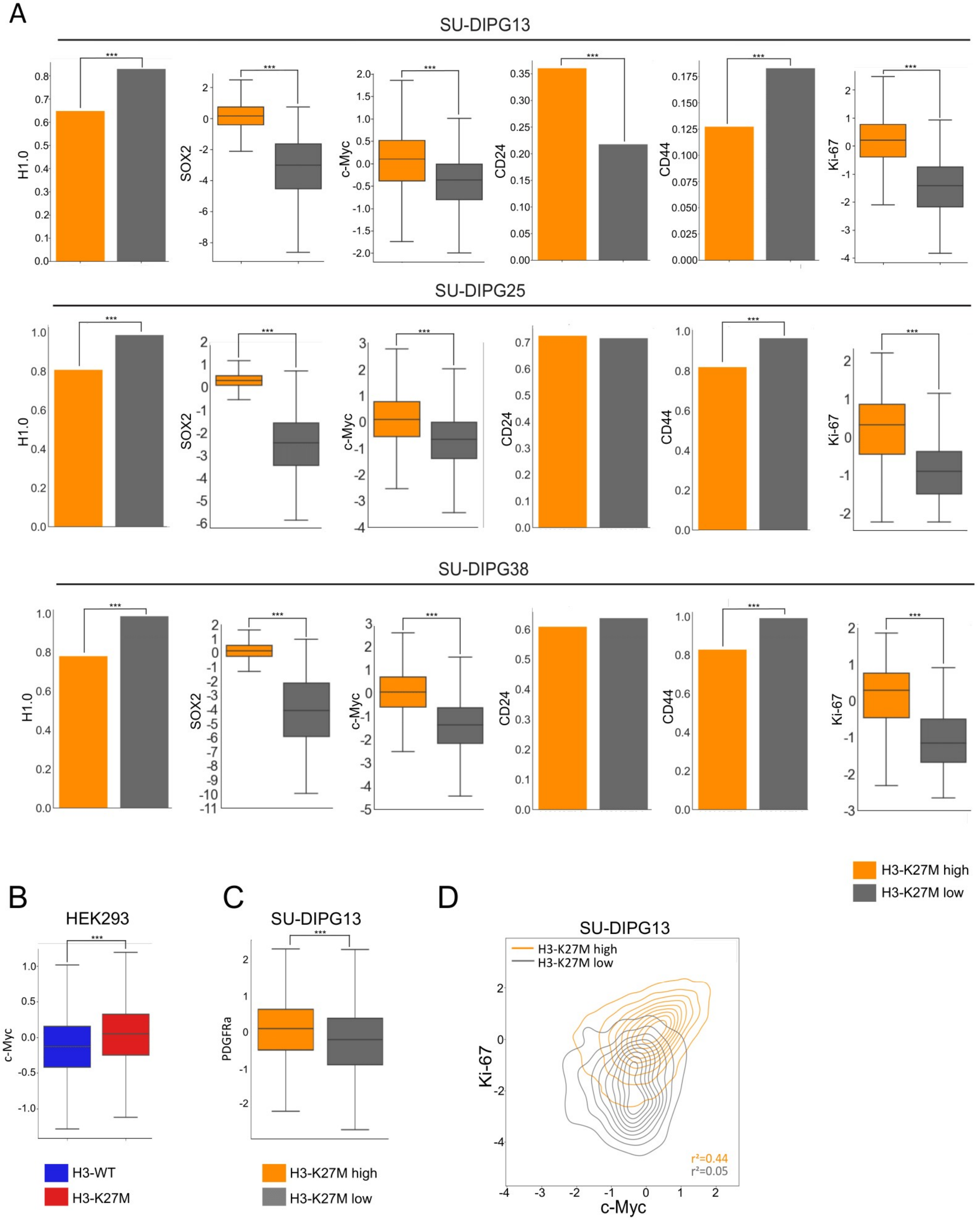
A
SU-DIPG13
SU-DIPG25
SU-DIPG38
H1.0
SOX2
c-Myc
CD24
CD44
Ki-67
H3-K27M high
H3-K27M low
B
HEK293
c-Myc
H3-WT
H3-K27M
C
SU-DIPG13
PDGFRa
H3-K27M high
H3-K27M low
D
SU-DIPG13
H3-K27M high
H3-K27M low
Ki-67
c-Myc
r²=0.44
r²=0.05

**Figure S5: The two epigenetic subpopulations show distinct proliferation capacity and expression of oncogenic and cancer stem-cells markers. Related to Figure 5.**

(**A**) Levels of the indicated oncogenic, cancer stem-cells and proliferation markers, as measured by CyTOF, for the H3-K27M-high and H3-K27M-low clusters in SU-DIPG13, SU-DIPG25 and SU-DIPG38. Left to right: Fraction of H1.0-positive cells, expression levels (transformed and scaled) of SOX2 and c-Myc, fraction of CD24 and CD44 positive cells, and expression levels of Ki-67. For H1.0, CD24 and CD44 fractions, p values were calculated by chi square test. For SOX2, c-Myc and Ki-67, p values were calculated by Welch's t-test. *** P value < 0.001. (**B**) Box plots of c-Myc expression levels, measured by CyTOF, in HEK293 cells expressing either WT or H3-K27M. P values was calculated by Welch's t-test. A single repeat was performed for this experiment. (**C**) PDGFRa expression levels in the two epigenetic clusters in SU-DIPG13, as measured by CyTOF. A single repeat was performed for this experiment, P values was calculated as in (B). (**D**) Contour plots of c-Myc versus Ki-67 in SU-DIPG13 H3-K27M-high (orange) and H3-K27M-low (gray) clusters. For panel composition and number of cells analyzed in all of these experiments, see Tables S2-3.

Fig. S6

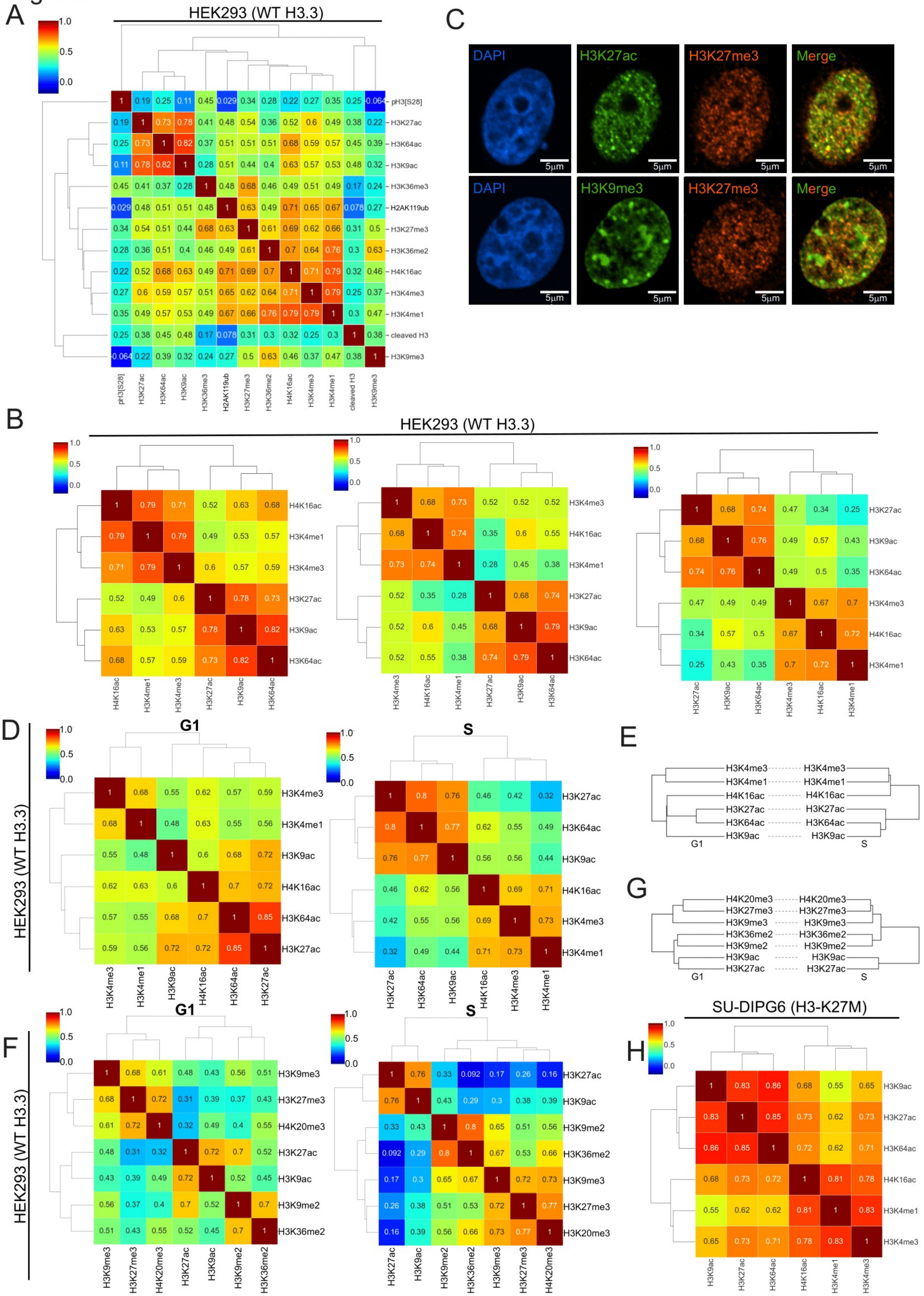
A
HEK293 (WT H3.3)
pH3[S28]
H3K27ac
H3K64ac
H3K9ac
H3K36me3
H2AK119ub
H3K27me3
H3K36me2
H4K16ac
H3K4me3
H3K4me1
cleaved H3
H3K9me3
C
DAPI
H3K27ac
H3K27me3
Merge
H3K9me3
5μm
B
HEK293 (WT H3.3)
D
HEK293 (WT H3.3)
G1
S
E
H3K4me3
H3K4me1
H4K16ac
H3K27ac
H3K64ac
H3K9ac
G
H4K20me3
H3K27me3
H3K9me3
H3K36me2
H3K9me2
H3K9ac
H3K27ac
F
HEK293 (WT H3.3)
G1
S
H
SU-DIPG6 (H3-K27M)

**Figure S6: Pairwise correlations of histone modifications. Related to Figure 6.**

**(A-B)** Pearson pairwise correlation between histone modifications measured by CyTOF (after transformation, scaling and normalization) in WT HEK293 cells. Shown is the coefficient of determination ($R^2$) heat map matrix of the following: (**A**) all histone modification in WT HEK293. **(B)** Biological repeats of WT HEK293, showing active histone modifications (similar to figure 6A). (**C**) Immunofluorescence staining of MCF7 cells with the indicated antibodies, DAPI was uswd for DNA labeling. H3K9me3 is localized to discrete foci in the nuclei, as opposed to H3K27me3 and H3K27ac that show a more dispersed pattern in the nucleoplasm. Scale = 5µm. (**D-G**) Pearson pairwise correlation between active (D and E) and repressive (F and G) histone modifications measured by CyTOF (after transformation, scaling and normalization) in HEK293 cells at G1 or S-phase. Cells were allocated to the indicated cell cycle phases based on cell cycle indicators included in the CyTOF panel. **(D, F)** The coefficient of determination ($R^2$) heat map matrix. **(E, G)** Dendograms of the indicated modifications in G1 versus S phase cells. The repressive modifications show similar clustering along the cell cycle. The active H4K16ac modification clusters with H3 acetylations during G1, yet in S-phase it clusters with H3K4me3 and H3K4me1. (**H**) Pearson pairwise correlation between active histone modifications measured by CyTOF (after transformation, scaling and normalization) in SU-DIPG6

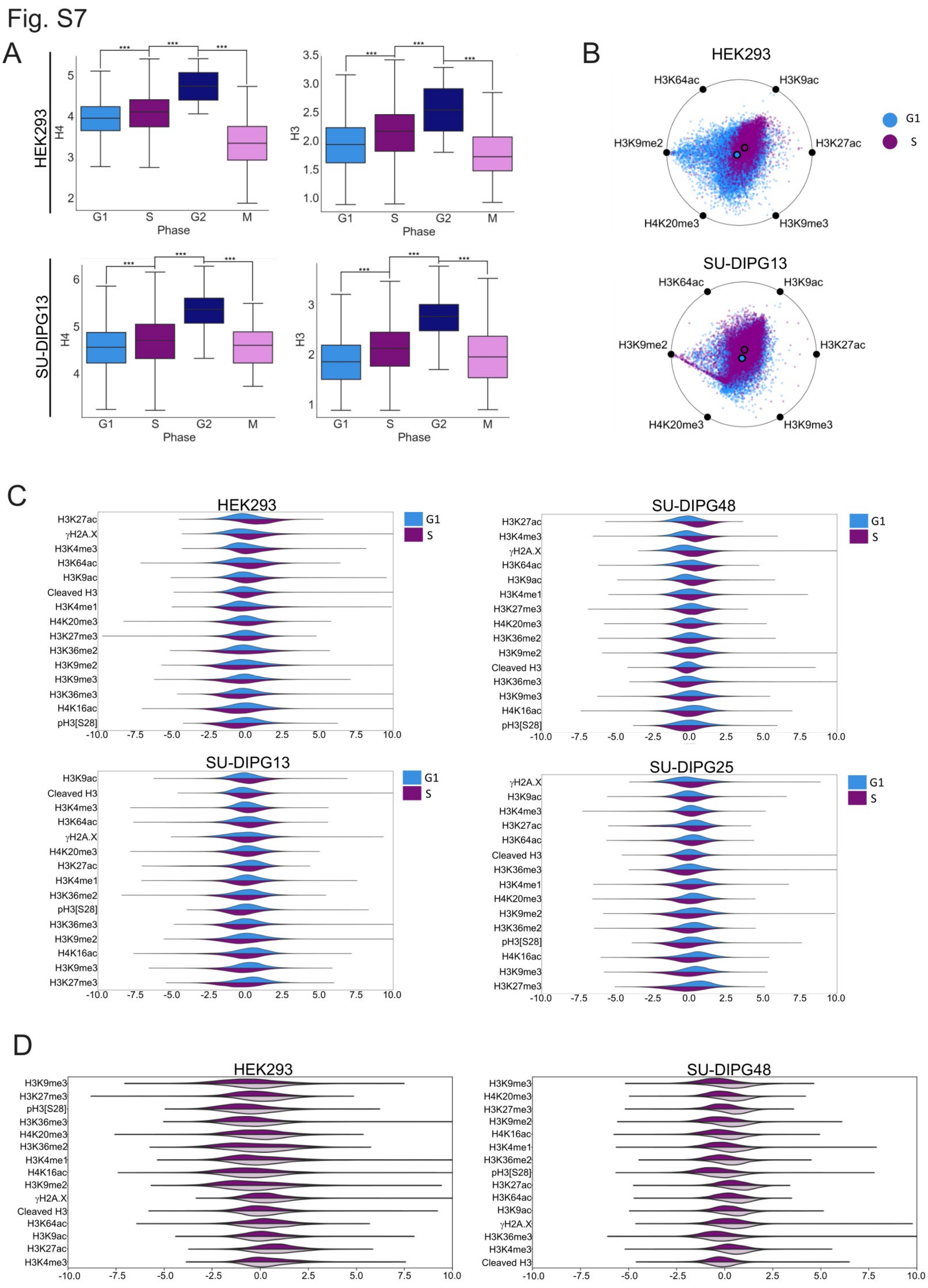
Fig. S7
A
HEK293
SU-DIPG13
H4
H3
Phase
G1
S
G2
M
***
B
HEK293
SU-DIPG13
H3K64ac
H3K9ac
H3K9me2
H3K27ac
H4K20me3
H3K9me3
G1
S
C
HEK293
SU-DIPG48
SU-DIPG13
SU-DIPG25
D
HEK293
SU-DIPG48

### Fig. S7 continued

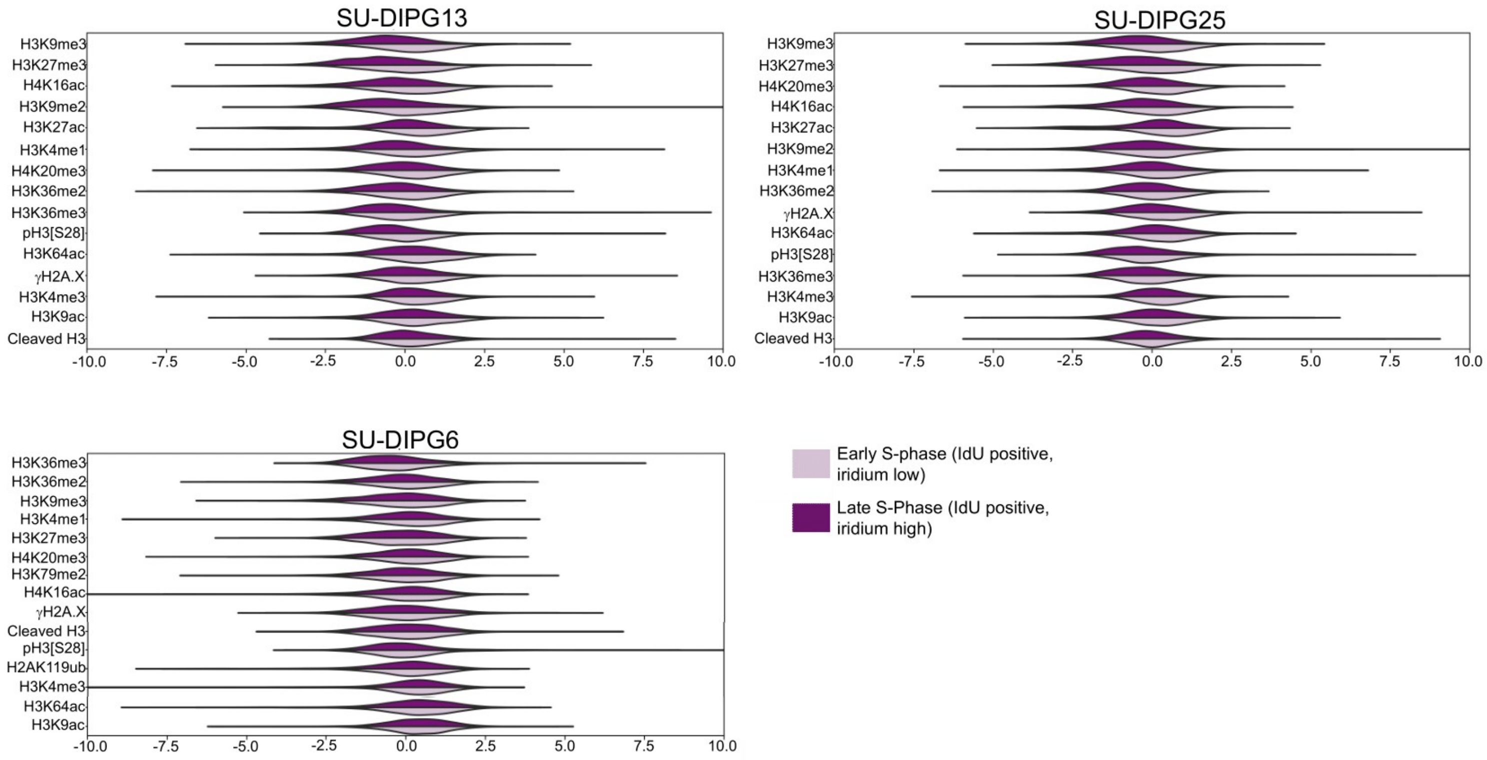

**Figure S7: High-dimensional single-cell analysis of epigenetic alterations during cell cycle. Related to Figure 7.**

(**A**) Box plots of the non-normalized scaled expression levels of the core histones in different phases of the cell cycle, in HEK293 and SU-DIG13 cells. P values were calculated for each phase in relation to the previous one by Welch's t-test. *** P value < 0.001. **(B)** Graphical representation of the epigenetic state of the G1 and S-phase populations of HEK293 and SU-DIPG13. (**C**) Data distribution of the epigenetic modifications, as measured by CyTOF, in cells at G1 and S phase, for the following cell lines: HEK293, SU-DIPG48, SU-DIPG13 and SU-DIPG25. (**D**) Data distribution of the epigenetic modifications, as measured by CyTOF, in cells at early and late S phase, for the following cell lines: HEK293, SU-DIPG48, SU-DIPG13 and SU-DIPG25 and SU-DIPG6.

**Table S1: Antibodies and metal allocations used in the CyTOF experiments. Related to STAR Methods.**

| Target | Metal | Name | Vendor | Cat # |
|---|---|---|---|---|
| H3 | 115In | Histone H3 Antibody | IonPath | 711501 |
| MBP | 140Ce | Anti-Myelin Basic Protein antibody [MBP101] | abcam | ab62631 |
| H3K36me3* | 141Pr | Tri-Methyl-Histone H3 (Lys36) (D5A7) XP® Rabbit mAb | CST | 4909 |
| GFAP | 143Nd | Anti-Cross GFAP (GA5) | Fluidigm | 3143022B |
| EZH2* | 144Nd | EZH2 (D2C9) XP® Rabbit mAb | CST | 5246 |
| H3K4me3* | 145nd | Tri-Methyl-Histone H3 (Lys4) (C42D8) Rabbit mAb | CST | 9751 |
| H3K79me2 | 146Nd | Anti-Histone H3 (di methyl K79) antibody - ChIP Grade | abcam | ab3594/ ab249947 |
| pHistone H2A.X | 147Sm | Anti-p-Histone H2A.X [Ser139] (JBW301) | Fluidigm | 3147016A |
| H3K36me2* | 149Sm | Di-Methyl-Histone H3 (Lys36) (C75H12) Rabbit mAb | CST | 2901 |
| SOX2 | 150Nd | Anti-Human SOX2 (O30-678) | Fluidigm | 3150019B |
| SIRT1 | 151Eu | Anti-SIRT1 antibody [19A7AB4] | abcam | ab110304 |
| H3K9me2 | 151Eu | Recombinant Anti-Histone H3 (di methyl K9) antibody [Y49] - BSA and Azide free | abcam | ab173325 |
| H4K16ac* | 152sm | Acetyl-Histone H4 (Lys16) (E2B8W) Rabbit mAb | CST | 13534 |
| H2Aub* | 153Eu | Rabbit monoclonal anti-ubiquityl-Histone H2A (Lys119) (clone D27C4) | CST | 8240 |
| H3K4me1* | 154sm | Mono-Methyl-Histone H3 (K4) (D1A9) XP(R) Rabbit mAb | CST | 5326S |
| H3.3 | 155Gd | Anti-Histone H3.3 antibody [EPR17899] - BSA and Azide free | abcam | ab208690 |

| | | | | |
|---|---|---|---|---|
| H3K64ac | 156Gd | Anti-Histone H3 (acetyl K64) antibody [EPR20713] – BSA and Azide free | abcam | ab251549 |
| BMI-1 | 157Gd | Anti-Bmi1 antibody [EPR22604-160] - BSA and Azide free | abcam | ab254475 |
| c-Myc | 158Gd | Anti-c-Myc antibody [Y69] - BSA and Azide free | abcam | ab168727 |
| H4 | 159Tb | Anti-Histone H4 antibody [mAbcam 31830] - ChIP Grade – BSA and Azide free | abcam | ab238663 |
| H3K27ac* | 160Gd | Acetyl-Histone H3 (Lys27) (D5E4) XP® Rabbit mAb | CST | 8173P |
| PDGFRa | 160Gd | Anti-Human PDGFRa (D13C6)-160Gd | Fluidigm | 3160007A |
| H4K20me3 | 161Dy | Anti-Histone H4 (tri methyl K20) antibody | abcam | ab9053 / ab239410 |
| DLL3 | 162Dy | Recombinant Anti-DLL3 antibody [EPR22592-18] - BSA and Azide free | abcam | ab255694 |
| Cleaved H3* | 163Dy | Rabbit monoclonal anti-cleaved-Histone H3 (Thr22) (clone D7J2K) | CST | 12576 |
| H3K9ac | 164Dy | Mouse monoclonal anti-acetyl-Histone H3 (Lys9) (clone 2G1F9) | Active Motif | 61663 |
| H1.0 | 165Ho | Anti-Histone H1.0 antibody [27] | abcam | ab11080 |
| H3K27ac | 165Ho | H3K27ac Monoclonal antibody | Thermo | MA5-23516 |
| CD24 | 166Er | Anti-Mouse CD24 | Fluidigm | 3166007B |
| H3K27me3 | 168Er | Mouse monoclonal anti-trimethyl-Histone H3 (Lys27) (clone MABI 0323) | Active Motif | 61017 |
| H3K27M | 169Tm | Recombinant Anti-Histone H3 (mutated K27M) antibody [EPR18340] - ChIP Grade – BSA and Azide free | abcam | ab240310 |
| H3K9me3* | 170Er | Tri-Methyl-Histone H3 (Lys9) (D4W1U) Rabbit mAb | CST | 13969S |

| | | | | |
|---|---|---|---|---|
| CD44 | 171Yb | Anti-Human/ Mouse CD44 (cancer stem cell marker) | Fluidigm | 3171003 |
| Ki-67 | 172Yb | Anti-Ki-67 (B56)-172Yb | Fluidigm | 3172024B |
| CXCR4 | 173Yb | Anti-Human CD184/CXCR4 (12G5) | Fluidigm | 3173001B |
| pH3[S28] | 175Lu | Anti-Human/ Mouse/ Rat pHistone H3 [Ser28] | Fluidigm | 3175012A |
| H1.3/4 | 176Yb | Anti-Histone H1.3 + Histone H1.4 antibody | abcam | ab61177 |

* Antibodies from CST arrived custom-made in a BSA-and-azide-free PBS solution, 1mg/ml concentration

**Table S2: Antibodies' panel composition per CyTOF experiment. Related to all figures.**

Panel composition (see CyTOF runs details below):

| Marker \ Run # | 1 | 2 | 3 | 4 | 5 | 6 | 7 | 8 | 9 | 10 | 11 | 12 | 13 | 14 | 15 |
|---|---|---|---|---|---|---|---|---|---|---|---|---|---|---|---|
| IdU-127I | | | | | v | v | | | v | v | | | | | |
| H3-115In | v | v | | v | v | v | v | v | v | v | v | v | v | v | v |
| MBP-140Ce | | | | | v | v | | | | | v | | | | |
| H3K36me3-141Pr | v | v | | v | v | v | v | v | v | v | v | | | | v |
| H3K27me2-142Nd | | | | | | | | | | | | v | v | v | |
| GFAP-143Nd | | | | | v | v | | | | | v | | | | |
| EZH2-144Nd | | | | | v | v | | | | | | | | | |
| H3K4me3-145nd | v | v | | v | v | v | v | v | v | v | v | | | | v |
| H3K79me2-146Nd | | | | | v | v | v | v | | | | | | | v |
| yH2A.X-147Sm | v | | | v | v | v | v | v | v | v | v | | | | v |
| H3K36me2-149Sm | v | v | v | v | v | v | v | v | v | v | v | v | v | v | v |
| pRb-150Nd | | | | | | | | | v | v | | | | | |
| SOX2-150Nd | | | | | v | v | v | v | | | v | | | | |
| H3K9me2-151Eu | | | | | | | v | v | v | v | v | v | | v | |
| SIRT1-151Eu | | | | | v | v | | | | | | | | | v |
| H4K16ac-152sm | v | v | v | v | v | v | v | v | v | v | v | v | v | v | v |
| CyclinB1-153Eu | | | | | | | | | v | v | | | | | |
| H2Aub-153Eu | v | v | | v | v | v | v | v | | | v | | v | v | v |
| H3K4me1-154sm | v | v | | v | v | v | v | v | v | v | v | | v | v | v |
| H3.3-155Gd | v | v | v | v | v | v | v | v | v | v | v | v | v | v | v |
| H3K64ac-156Gd | v | v | | v | v | v | v | v | v | v | v | v | v | v | v |
| BMI-1-157Gd | | | v | | v | v | | | | | | | | | |
| c-Myc-158Gd | | | v | | v | v | v | v | v | v | v | | | | |
| H4-159Tb | v | v | v | v | v | v | v | v | v | v | v | v | v | v | v |
| H3K27ac-160Gd | v | v | v | v | v | | v | v | v | v | v | v | | | v |
| PDGFRa-160Gd | | | | | | v | | | | | | | | | |
| H4K20me3-161Dy | | | | | v | v | v | v | v | v | v | | | | v |
| DLL3-162Dy | | | | | v | v | | | | | | | | | |
| Cleaved H3-163Dy | v | v | v | v | v | v | v | v | v | v | v | v | v | v | v |

| | | | | | | | | | | | | | | | |
|---|---|---|---|---|---|---|---|---|---|---|---|---|---|---|---|
| **H3K9ac-164Dy** | v | v | | v | v | v | v | v | v | v | v | | | | v |
| **H3K27ac-165Ho** | | | | | | v | | | | | | | | | |
| **H1.0-165Ho** | | | | | v | | v | | | | | | | | v |
| **CD24-166Er** | | | | | v | v | v | v | v | | v | | | | |
| **H3K27me3-168Er** | v | v | v | v | v | v | v | v | v | v | v | v | v | v | v |
| **H3K27M-169Tm** | v | v | v | v | v | v | v | v | v | | v | v | v | v | |
| **H3K9me3-170Er** | v | v | v | v | v | v | v | v | v | v | v | v | v | v | v |
| **CD44-171Yb** | | | | | v | v | v | v | | v | v | | | | |
| **Ki-67-172Yb** | | | | | v | v | v | v | v | v | v | | | | |
| **CXCR4-173Yb** | | | | | v | | | v | | v | v | | | | |
| **pH3[S28]-175Lu** | v | v | v | v | v | v | v | v | v | v | v | v | v | v | |
| **H1.3/4-176Yb** | | | | | v | | | | | | | | | | |

Experiment according to CyTOF runs:

| **#** | **Samples and related figures** |
|---|---|
| **1** | HEK293 K27M #1 10d induction (1B,C and S1E-G and 6E)<br>HEK293 WT #1 10d induction (1B,C and S1E,F, S6AB) |
| **2** | HEK293 K27M #2 4d induction (S1E,F)<br>HEK293 WT #2 4d induction (S1E,F, and S6B)<br>HEK293 K27M #3 7d induction (S1E,F)<br>HEK293 WT #3 7d induction (S1E,F and S6B) |
| **3** | HEK293 dynamics WT (3A-D, S3A and S5B)<br>HEK293 dynamics K27M 8,16,48 & 96 hrs (3A-D and S3A) |
| **4** | SU-DIPG48 #1 (2A)<br>SU-DIPG13 #1 (2A-D, S2B-F, and 4F) |
| **5** | SU-DIPG6 (4A,C,D, S4C,E,J, 5A,B,D-G, S5D, 6SH, 7F,G and S7D)<br>SU-DIPG13 #2 (4A,C,F, S4D,E, S4D,E,J, 5A and 6F) |
| **6** | SU-DIPG13 alternative panel with PDGFRa (S5C) |
| **7** | SJ-HGGX39 (4B)<br>SU-DIPG38 (4A, S4B,E, 5A and S5A)<br>SU-DIPG25 #1 (4A,B,E, S4E, 5A and S5A) |

| | |
|---|---|
| 8 | SU-DIPG36 K27M (S2G, 3G and 3SB)<br>SU-DIPG36 KO (S2G, 3G and 3SB)<br>BT245 K27M (S2H and 3G)<br>BT245 KO (S2H and 3G) |
| 9 | SU-DIPG48 #2 cell cycle (S2A, 4B, 6B,D, 7A-G and S7C,D)<br>SU-DIPG13 #3 cell cycle (S2A, 4B,C,F, 6A,C, 7B,F,G and S7A-D)<br>SU-DIPG25 #2 cell cycle (4C,E, S4A,F-I, 6G, 7B,F,G and S7C,D)<br>SU-DIPG25 #3 cell cycle (-) |
| 10 | HEK293 cell cycle (S6D-F, 7B,F,G and S7A-D) |
| 11 | SU-DIPG13 #4 (vorino 0) (1SB, 4F and 6H,I)<br>SU-DIPG13 (vorino 12,24,36 hrs) (6H,I)<br>SU-DIPG13 (vorino 48 hrs) (S1B and 6H,I) |
| 12 | SJ-HGGX39V WT H3 (S1C)<br>SJ-HGGX39V Dynamics K27M dox 8,16,48 & 96 hrs (3E,F and S3C,D) |
| 13 | SU-DIPG13 OE (4G)<br>SU-DIPG13 OE uninduced (4G) |
| 14 | SU-DIPG13 shControl (4H and S4K,L)<br>SU-DIPG13 shK27M (4H) |
| 15 | HEK293 siControl (S1A,C)<br>HEK293 siKMT2D (s1A)<br>HEK293 siKMT2A (S1A)<br>HEK293 siMOF (S1C) |

**Table S3: Number of cell analyzed per CyTOF experiment. Related to all figures.**

| Cell line and repeat number | Relate to Figure | Total live single cells analyzed | H3-K27M-High Cells (%) | H3-K27M-Low Cells (%) |
|---|---|---|---|---|
| SU-DIPG13 #1 | 2A-D, S2B-F, and 4F | 12081 | 54.4 | 45.6 |
| SU-DIPG13 #2 | 4A,C,F, S4D,E, S4D,E,J, 5A and 6F | 14979 | 93.7 | 6.3 |
| SU-DIPG13 #3 (cell cycle) | S2A, 4B,C,F, 6A,C, 7B,F,G and S7A-D | 27149 | 90.9 | 9.1 |
| SU-DIPG13 #4 (vorino 0) | 1SB, 4F and 6H,I | 62323 | 87.6 | 12.4 |
| SU-DIPG13 4 (vorino 12hr) | 6H,I | 57702 | N/A | N/A |
| SU-DIPG13 4 (vorino 24hr) | 6H,I | 49863 | N/A | N/A |
| SU-DIPG13 4 (vorino 36hr) | 6H,I | 52400 | N/A | N/A |
| SU-DIPG13 4 (vorino 48hr) | S1B and 6H,I | 46966 | N/A | N/A |
| SU-DIPG13 OE | 4G | 29042 | 97.7 | 2.3 |
| SU-DIPG13 OE uninduced | 4G | 25210 | 94.8 | 5.2 |
| SU-DIPG13 alternative panel with PDGFRa | S5C | 12775 | N/A | N/A |
| SJ-HGGX39V WT H3 | S1C | 138173 | N/A | N/A |
| SJ-HGGX39V K27M dox 8hr | 3E,F and S3C,D | 175756 | N/A | N/A |
| SJ-HGGX39V K27M dox 16hr | 3E,F and S3C,D | 140773 | N/A | N/A |
| SJ-HGGX39V K27M dox 48hr | 3E,F and S3C,D | 140382 | N/A | N/A |
| SJ-HGGX39V K27M dox 96hr | S1C, 3E,F and S3C,D | 142921 | N/A | N/A |
| SU-DIPG25 #1 | 4A,B,E, S4E, 5A and S5A. | 70555 | 89.4 | 10.6 |

| | | | | |
|---|---|---|---|---|
| SU-DIPG25 #2 (cell cycle) | 4C,E, S4A,F-I, 6G, 7B,F,G and S7C,D | 21549 | 81.7 | 18.3 |
| SU-DIPG25 #3 (cell cycle) | - | 18372 | 84.3 | 15.7 |
| SU-DIPG6 | 4A,C,D, S4C,E,J, 5A,B,D-G, S5D, 6SH, 7F,G and S7D | 19983 | 89.8 | 10.2 |
| SU-DIPG38 | 4A, S4B,E, 5A and S5A | 57616 | 77 | 23 |
| SU-DIPG48 #1 | 2A | 8190 | N/A | N/A |
| SU-DIPG48 #2 (cell cycle) | S2A, 4B, 6B,D, 7A-G and S7C,D | 23257 | N/A | N/A |
| SJ-HGGX39 | 4B | 130745 | N/A | N/A |
| SU-DIPG13 shControl | 4H and S4K,L | 11008 | See Fig4H | See Fig4H |
| SU-DIPG13 shK27M | 4H | 10248 | See Fig4H | See Fig4H |
| SU-DIPG36 K27M | S2G, 3G and 3SB | 4209 | N/A | N/A |
| SU-DIPG36 KO | S2G, 3G and 3SB | 2711 | N/A | N/A |
| BT245 K27M | S2H and 3G | 1034 | N/A | N/A |
| BT245 KO | S2H and 3G | 2828 | N/A | N/A |
| HEK293 (cell cycle) | S6D-F, 7B,F,G and S7A-D | 34606 | N/A | N/A |
| HEK293 K27M #1 10d induction | 1B,C and S1E-G and 6E | 22147 | N/A | N/A |
| HEK293 WT #1 10d induction | 1B,C and S1E,F, S6AB | 12760 | N/A | |
| HEK293 K27M #2 4d induction | S1E,F | 31422 | N/A | N/A |

| | | | | |
|---|---|---|---|---|
| HEK293 WT #2 4d induction | S1E,F, and S6B | 37379 | N/A | N/A |
| HEK293 K27M #3 7d induction | S1E,F | 23127 | N/A | N/A |
| HEK293 WT #3 7d induction | S1E,F and S6B | 22266 | N/A | N/A |
| HEK293 dynamics WT | 3A-D, S3A and S5B | 84118 | N/A | N/A |
| HEK293 dynamics 8hr | 3A-D and S3A | 59774 | N/A | N/A |
| HEK293 dynamics 16hr | 3A-D and S3A | 82085 | N/A | N/A |
| HEK293 dynamics 48hr | 3A-D and S3A | 72638 | N/A | N/A |
| HEK293 dynamics 96hr | 3A-D and S3A | 73053 | N/A | N/A |
| HEK293 siControl | S1A,C | 58634 | N/A | N/A |
| HEK293 siKMT2D | S1A | 59449 | N/A | N/A |
| HEK293 siKMT2A | S1A | 58710 | N/A | N/A |
| HEK293 siMOF | S1C | 75287 | N/A | N/A |
